# A genomic platform for epidemiological surveillance and vaccine antigen discovery using long-read amplicon sequencing

**DOI:** 10.1101/2022.06.08.495295

**Authors:** David Fernando Plaza, Julia Zerebinski, Ioanna Broumou, Maximilian Julius Lautenbach, Billy Ngasala, Christopher Sundling, Anna Färnert

## Abstract

Many vaccine candidate proteins are under strong selective pressure to diversify in terms of antigenicity. We present a sequencing and data analysis platform for epidemiological surveillance and discovery of indel-rich vaccine antigens by long-read circular consensus sequencing (CCS) in multiclonal pathogen isolates. Our platform uses 40 PCR primers to asymmetrically barcode and identify multiclonal infections in pools of up to 384 samples. We validated the method using 235 mock infections combining 10 synthetic variants of the indel-rich gene merozoite surface protein 2 of *Plasmodium falciparum* at different concentrations and infection complexities, as well as 95 isolates from *P. falciparum*-infected residents of Nyamisati, Tanzania. We also constructed a fully automated analysis pipeline that streamlines the processing and interpretation of epidemiological and antigenic diversity data from demultiplexed FASTQ files. This platform can be easily adapted to other polymorphic antigens of interest in *Plasmodium* and other human pathogens.

---

There were 241 million cases of malaria and 627 thousand deaths from the disease in 2020^1^. An effective vaccine against malaria is a cost-effective and urgently needed strategy to reduce the disease burden and potentially eradicate malaria in the future^2^. RTS,S/AS01 (Mosquirix™) is the first malaria vaccine to be recently endorsed by the World Health Organization for use in children^3^. Nevertheless, the protection conferred by RTS,S/AS01 is strain-specific^4^, therefore limiting the effectiveness of this vaccine in regions where *P. falciparum* circulates with a high genetic and antigenic diversity^5^. New approaches to survey the pool of genetic diversity in highly polymorphic antigens from *P. falciparum* and other pathogens will accelerate the development of strain-transcending vaccines.

Merozoite surface protein 2 (*msp2*) is highly expressed in the blood stage of *P. falciparum*^6^. The protein has two highly conserved domains in the N and the C termini flanking a highly polymorphic central region. The locus is a hotspot for indels of up to 300bp^7^, making its sequencing and assembly, especially for multiclonal infections, a challenging task using short-read sequencing^8^. Insertions, deletions or tandem repeats larger than the read lengths in short-read sequencing represent another significant mapping and assembly challenge.

Here, we present a multiplex long-read amplicon sequencing platform for the accurate genotyping of indel-rich antigens using the malaria vaccine candidate MSP2 as a proof of concept. The use of this circular consensus sequencing (CCS) technology to obtain single-base resolution data on polymorphic antigens will facilitate the implementation of a comprehensive genomic surveillance strategy for infectious diseases caused by highly diverse pathogens and will allow the in-depth structural and antigenic characterization of vaccine candidates capable of eliciting strain-transcending immunity.

## Results

### CCS on pools of amplicons provides insights into complex infections with high sequence accuracy

Combining nested PCR, asymmetric barcoding of amplicons and CCS, we developed a platform for the genotyping by sequencing of loci rich in structural variants, in up to 384 pooled controls and clinical isolates (Fig. 1). With an average read length of 28300 bp and a mean insert size of 862 bp, every *msp2* amplicon was sequenced 32 times on average, generating circular consensus reads where most sequencing errors (previously estimated to be one error every 13048 bp in CCS reads^9^) are properly amended. We observed no correlation between template concentration in a series of mock infection controls including single *msp2* variants and the number of reads resulting from those samples (Supplementary Fig. 1). Similarly, there was no correlation between sample parasite density (parasites/µl) and the number of CCS reads for microscopy-positive samples in the Tanzania cohort (Supplementary Fig. 2).

**Fig 1.**
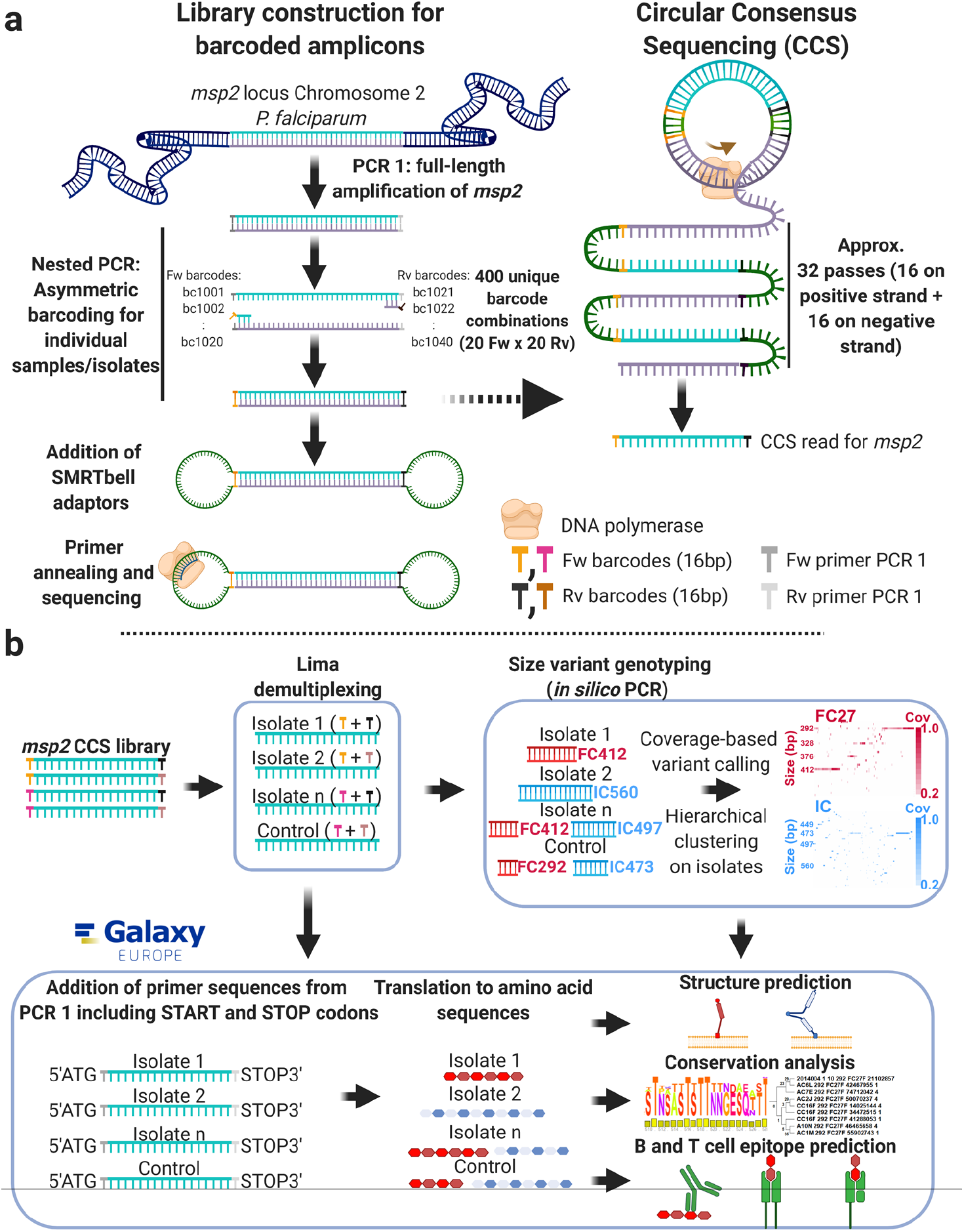
Library construction and analysis pipeline for *msp2* genotyping by circular consensus sequencing. **a**, Full-length *msp2* was amplified in a first round of PCR that used primers annealing to the conserved 5’ and 3’ ends of the *msp2* gene. In the nested PCR that followed, *msp2* amplicons were asymmetrically barcoded using primers from the conserved 5’ and 3’ regions of the gene but annealing directly downstream and upstream from the annealing sites for the forward and reverse oligos, respectively, used in the first PCR reaction. Asymmetric amplicon barcoding allowed for the accurate pooling and computational identification of up to 384 individual samples using a combination of 40 different barcodes. Thereafter, SMRTbell adaptors were coupled to the barcoded amplicons resulting in single-stranded circularized constructs containing both strands of *msp2*. Circularized *msp2* amplicons were sequenced in 32 polymerase passes (16 on each strand) and the resulting sequences for each amplicon were aligned to produce circular consensus reads. **b**, Circular consensus sequences (CCSs) were de-multiplexed with Lima and a combined FASTA file was created with all these indexed CCSs reads. Size variant genotyping was done by *in silico* PCR on this FASTA file using the IC1 and FC27 primer sequences designed by Snounou *et al*, 1999, in addition to new primers designed for this study as queries in a Blastn search against the dataset. Isolates and size variants were hierarchically clustered and visualized in two sequencing coverage heatmaps for IC1 and FC27. In addition, CCS reads were bioinformatically coupled to the primers from the first PCR to construct isolate phylogenies or be translated into amino acid sequences to allow the prediction of structural elements, the study of sequence conservation, as well as the presence of B and T cell epitopes.

### Variant calling specificity and sensitivity can be estimated from synthetic mock infection controls

Size and sequence variant calling for clinical isolates was based on the accuracy of variant detection in synthetic mock infection controls. These controls corresponded to 235 single or mixed serial dilutions including various molar ratios of synthetic reference sequences for *msp2* HB3 (LR131339 REGION: 250760..251530), CD01 (PfCD01_020011700), Dd2 (PfDd2_020009600), SN01 (PfSN01_020009800), KE01 (PfKE01_020009300), SD01 (PfSD01_020012300), GN01 (PfGN01_020012100), 7G8 (Pf7G8_020011500), GB4 (PfGB4_020009800) and 3D7 (PF3D7_0206800), that were barcoded separately and pooled together before adaptor ligation and sequencing (Supplementary Table 1). Serial dilution of single-variant mock infections showed that the correct size (Fig. 2a) and sequence (Fig. 2b) variant for reference alleles can be detected with read coverages higher than 50% at concentrations of 10 copies/μl for all the variants and down to 1 copy/μl for 6 out of 10. We observed that all samples with barcode bc1034, including the 100 copies/μl dilution for variants SD01, GN01, 7G8, GB4, CD01, Dd2, SN01 and KE01, did not yield reads above background levels (Supplementary Fig. 3) and were therefore not included in the sensitivity analysis.

**Fig 2.**
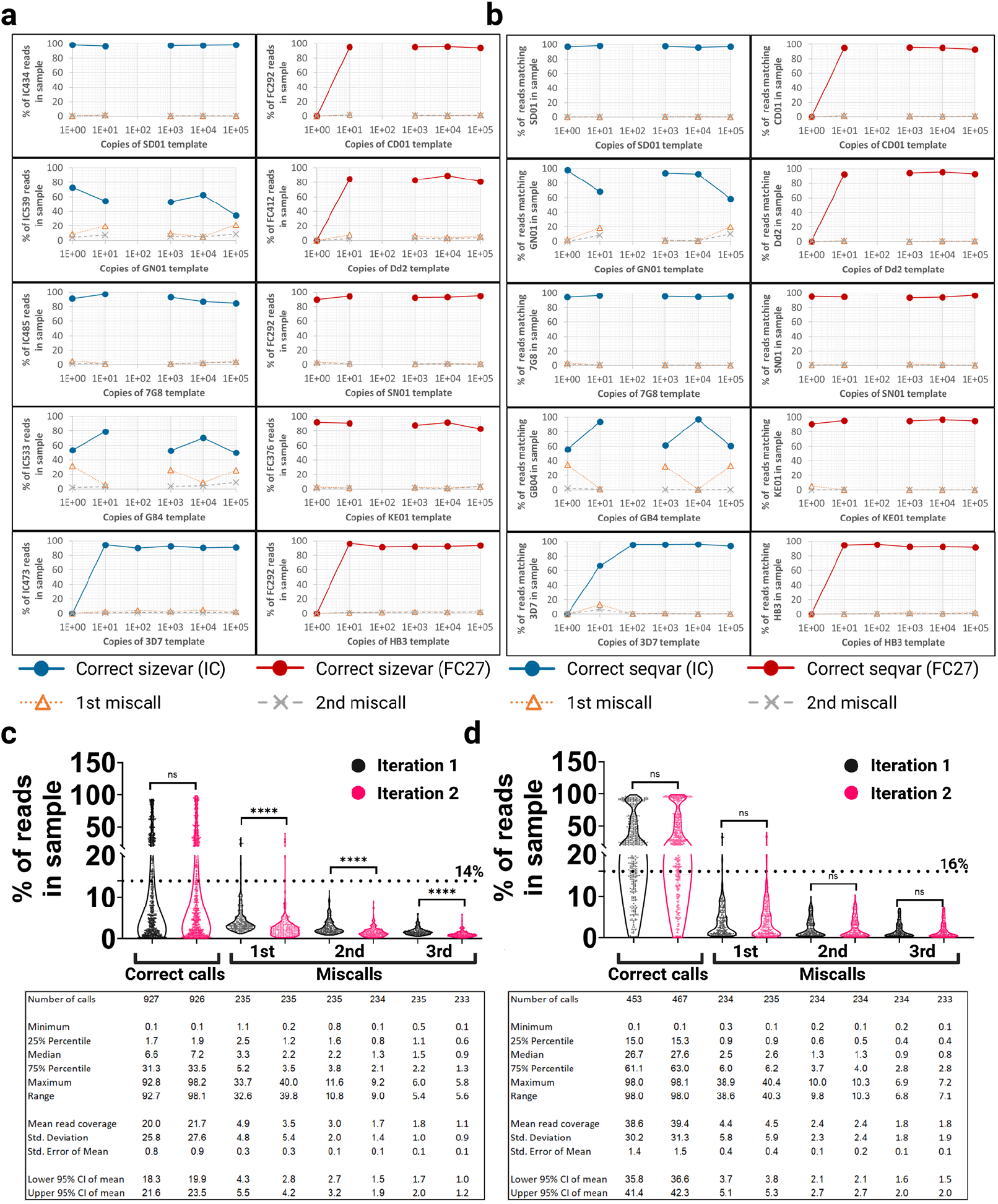
Sensitivity and specificity in the calling of size and sequence variants, as well as variant calling thresholds can be established using mock infections with synthetic constructs of *msp2*. Serial dilutions of pure synthetic *msp2* from the reference lines SD01 (IC434), GN01 (IC539), 7G8 (IC485), GB4 (IC533), 3D7 (IC473), CD01 (FC292), Dd2 (FC412), SN01 (FC292), KE01 (FC376) and HB3 (FC292) were prepared and individually barcoded (each step in the dilution series with a unique barcoding combination), pooled together and sequenced. Dataset noise was measured as the read coverage for the most frequent miscalls in each sample. The correct **a**, size and **b**, sequence variants were called for a range of template concentrations (number of template copies) in single-clone mock infections. Coverage for the correct variant (Correct sizevar or Correct seqvar), the most common variant miscall (1^st^ miscall) and the second most common variant miscall (2^nd^ miscall) in the IC1 (in blue) and FC27 (in red) families of *msp2* are shown. Percentage of reads per sample for **c**, size and **d**, sequence variant correct calls and miscalls present in a set of 235 synthetic mock infections of different complexity (from 1 to 10 clones per infection mixed in different molar ratios to emulate possible infection scenarios found in clinical isolates). Read coverage was measured in the first and second iterations of the analysis workflow, showing that the noise reduction module introduced between the two iterations significantly reduces the percentage of reads in the sample assigned to the three most frequent size variant miscalls (Mann-Whitney pairwise comparison, **** P value < 0.0001). Mean and SD are shown. Based on these, thresholds for variant calling were set to the mean coverage of the most common miscall plus two standard deviations, being these 14% and 16% for size and sequence variants (shown as dotted lines in **c**, and **d**,), respectively.

Quantification of left-over noise after the noise-reduction module (see below) revealed that the mean read coverage for the most common size variant miscall in 235 mock infections was 3.5% of reads (SD 5.4, 95% CI 2.8-4.2%) (Fig. 2c). A downstream analysis of sequence variants for the same pool of mock infections showed that the mean read coverage for the most common sequence variant miscall was 4.5% of the reads (SD 5.9, 95% CI 3.8-5.3%) (Fig. 2d). The mean read coverage for size variant miscalls at different molar ratios and dilutions in the synthetic mock infections (Fig. 2c) allowed us to setup a variant calling threshold of 14% (the mean read coverage of the most prevalent variant miscall in all the mock infections plus 2 standard deviations (SD)). Similarly, a coverage threshold was defined in the calling of nucleotide sequence variants (Fig. 2d) equivalent to 16% or the mean coverage of the most common sequence miscall in the synthetic controls plus 2 SD. For the same 235 mock infections, a total of 1140 correct size variant calls were expected; nevertheless, 926 (81.2%) of those were detected in the dataset with a total of 360 above the noise threshold (31.2% of the expected, false negative rate (FNR): 0.688). Furthermore, 7407 size variant miscalls were detected, from which only 9 had read coverages above or equal to the established 14% threshold (False positive rate (FPR): 0.001). As for sequence variants, 1312 correct calls were expected in the 235 mock infections. From these, 467 (35.6%) were detected with 342 above the noise threshold (26.1% of the expected, FNR: 0.739). In addition, 8082 sequence variant miscalls were detected and only 8 of those showed read coverages above the corresponding variant calling cutoff of 16% (FPR: 0.001). Finally, size and sequence variant calling thresholds at FPRs of 0.01 and 0.05 were calculated (Supplementary Table 2). The selection of FPR significantly affects the number of variants that can be detected in individual isolates, with less stringent FPRs (lower variant calling thresholds) resulting in a closer match between the number of variants (clones in a clinical isolate, else referred as multiplicity of infection (MOI)) expected and observed, however compromising the sequence accuracy of the variants called (Supplementary >Fig. 4).

### Size variant genotyping for clinical isolates can be simulated by in silico PCR on CCS reads

Even though sequencing with single nucleotide resolution is the optimal tool to assess isolate complexity and relatedness in molecular epidemiological surveys ^10, 11^, most of the available genetic diversity data for *P. falciparum* relies on the genotyping of size variants in *msp2, msp1* and *glurp*^12^. *In silico* PCR on CCS reads with the widely used family-specific (IC1 and FC27) primers for *msp2*^13^ was applied to call size variants on isolates which could then be compared to data previously generated by nested PCR (using the same primers) and analysis of fragment sizes by capillary electrophoresis (CE)^14^. Fully sequenced and annotated genomes for a number *P. falciparum* strains became available (PlasmoDB^15^) in the years following the original *msp2* genotyping publication^13^ and we used these sequences to design forward oligos that incorporate new base substitutions (Supplementary Fig. 5 and Supplementary Table 3).

*In silico* PCR on the library of synthetic mock infections using the original and the newly designed primers followed by size variant calling showed that the eight size variants expected (three different sequence variants correspond to FC292) were detected. In addition, a clinical isolate corresponding to an FC316 variant and spiked into some of the mock infection samples was found in 10 samples at an FPRs of 0.001 (Fig. 3a). Only three miscalls were detected in 5 out of the 321 (1.6%) mock infections analyzed using the same FPR. Lastly, *in silico* PCR for size variant genotyping in clinical isolates that were sequenced in a separate library, showed that 52 (54.7%) out of 95 isolates were IC1-positive (Fig. 3b) while 59 (62.1%) were FC27-positive (Fig. 3c). Sixteen samples (16.8%) were found to be double positive for IC1 and FC27.

**Fig 3.**
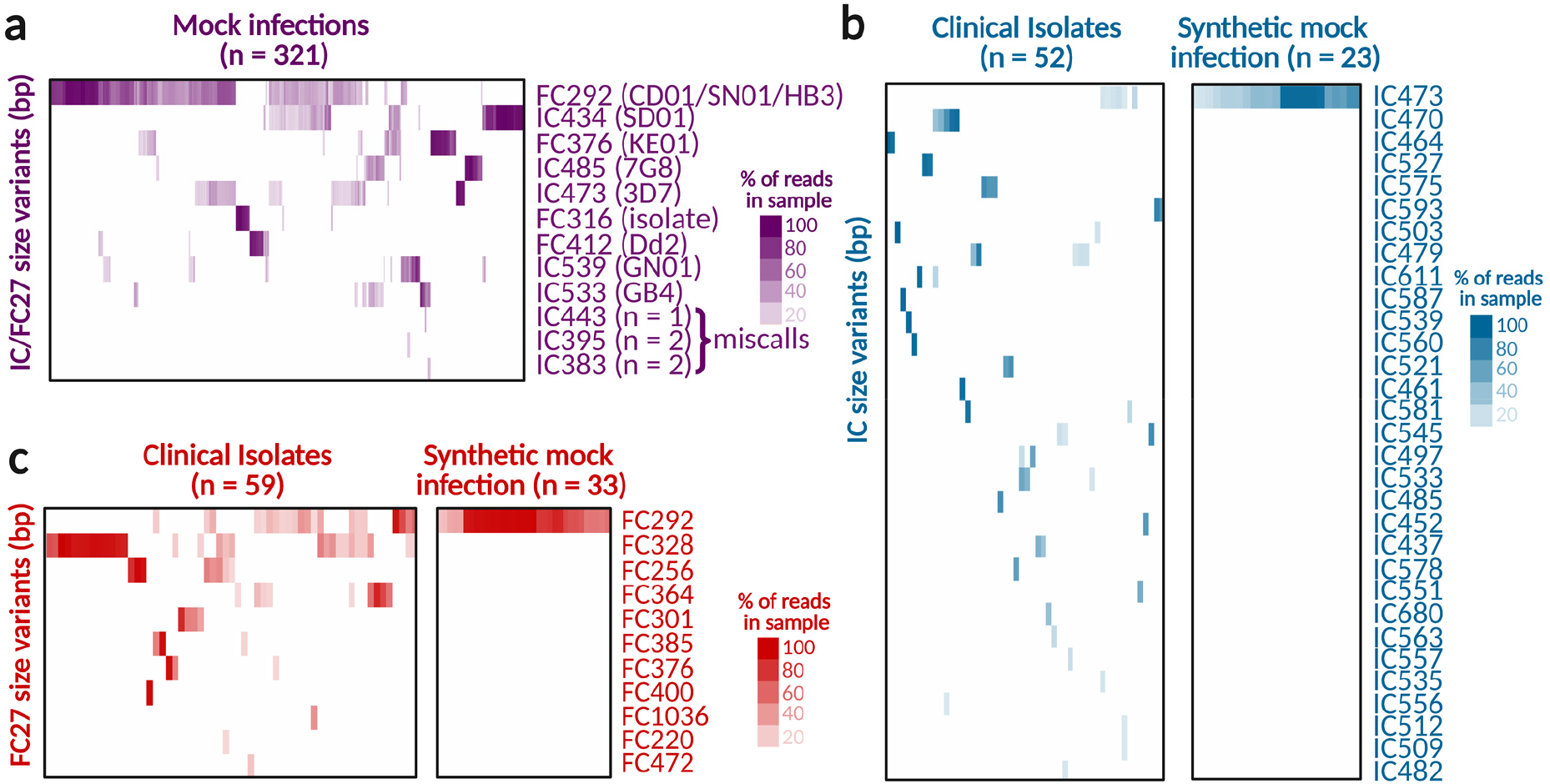
Size variant genotyping simulated by *in silico* PCR on CCS reads from synthetic mock infections and clinical isolates. **a**, Variant calling for 321 mock infection controls. Mock infections were prepared with 10 synthetic gene variants of *msp2* (in parenthesis) in concentrations of 1 to 100000 copies per reaction. Infections ranged in complexity from 1 to 10 variants per mixture. One clinical isolate (FC316) was spiked-in and detected in 11 samples included in the pool. IC443, IC395 and IC383 are miscalls present in 5 out of 321 mock infections (1.6%). **b**, IC1 and **c**, FC27 size variant genotyping for 95 clinical isolates collected in Nyamisati, Tanzania, in 1999 and 2016. Size variant calling is shown for synthetic mock infections used as positive controls in the CCS libraries where the isolates were sequenced and including a mix of one IC1 (3D7, IC473) and one FC27 (HB3, FC292) variant. Isolates, mock infections and size variants were grouped by complete hierarchical clustering using Euclidian as the distance metric.

Even though amplification conditions vary widely in the PCR programs for size variant genotyping by CE and the barcoding of *msp2* amplicons, we decided to compare the specific size variants being called by CE and CCS on 24 *msp2*-positive samples from Tanzania that were randomly selected. A 7bp plus or minus window in the CE calls was used to account for the size variant binning required when using this method^14^. We found that for 15 out of 24 *msp2*-positive samples (62.5%, Supplementary Table 4) there was at least one match in the size variants called by the two methods with high relative fluorescence units (RFU>300) and read coverage values (≥14% of reads in the sample).

### Isolate-derived msp2 sequences allow the analysis of antigen structural conservation and the construction of isolate phylogenies

Finally, isolate phylogenies, analysis of B-cell epitope conservation and prediction of T cell epitopes were computed for the same 95 *P. falciparum* isolates. Conservation analysis of unique MSP2 variants of *P. falciparum* in the IC1 (n = 37) and FC27 (n = 34) families for isolates from Tanzania shows that there are well defined regions which are highly conserved in the two families (Fig. 4a) and that could be targeted by vaccine-elicited antibodies to confer strain-transcending protection. We used the consensus sequences derived from the multiple alignment Fourier transform (MAFFT) of the MSP2 variants present in the isolates to map linear epitopes bound by well-characterized monoclonal antibodies (mAbs) raised against the 3D7 and FC27 variants^16^. We observed that epitopes present in conserved regions (6D8, 1F7/6C9/9H4 and 4D11/9G8) can also be present in the variable regions of some of the clinical isolates sequenced (Fig. 4a). Furthermore, sequences of a polymorphic locus like *msp2* are instrumental in the construction of phylogenies describing the degree of relatedness between different *P. falciparum* isolates. We show that *msp2* sequences from clinical isolates can be used to construct phylogenies for the two families of the gene (Fig. 4b and Fig. 4c).

**Fig 4.**
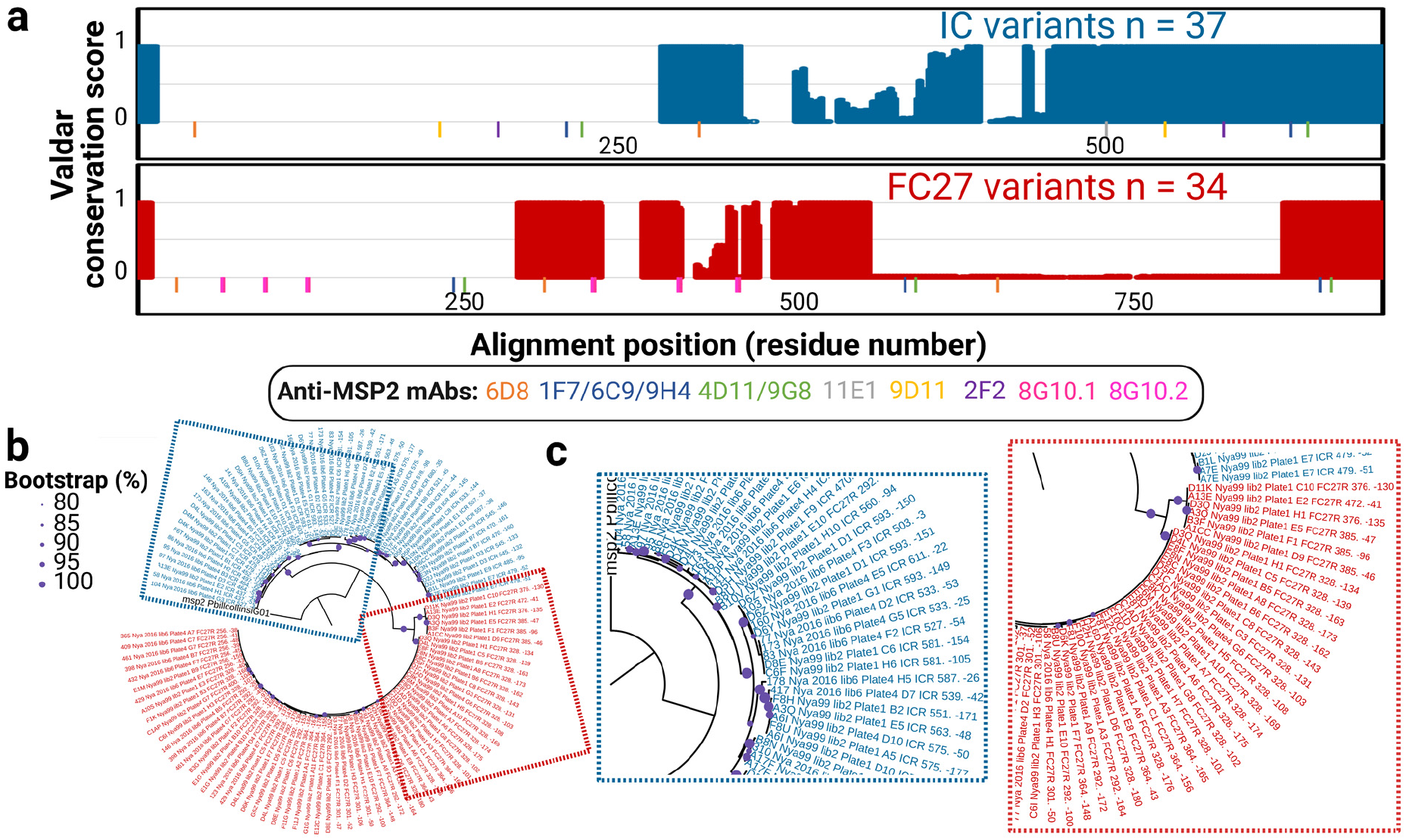
Conservation of MSP2 protein sequences and phylogenetic analysis of malaria isolates based on *msp2*. Amino acid and nucleotide sequences were aligned with MAFFT. **a**, Valdar conservation scores were plotted for every residue position in the alignment for the IC1 and FC27 families. MSP2 regions potentially targeted by broadly-neutralizing (strain-transcending) antibodies show high Valdar conservation scores. The location of epitopes recognized by previously characterized mouse monoclonal antibodies is shown under each conservation plot. **b**, Maximum likelihood phylogenetic tree for 51 and 62 unique clones of the IC1 (in blue) and FC27 (in red) families, respectively, found in isolates collected in Tanzania in 1999 and 2016. Tree branches supported by 80% or more of the bootstrapping replicates are marked with purple circles. The tree was rooted using the ortholog of *msp2* present in the species *P. billcollinsi* (*Laverania* subgenus). **c**, The detailed structure of the trees is shown in the dashed frame panels.

### Non-redundant T cell epitopes with wide human population coverage were detected in the MSP2 variants from a set of clinical isolates

To explore how sequence diversity could be affecting the immunogenicity of different antigen variants, we computed binding predictions for all the MSP2 15-mer peptides from the same Tanzania isolates against the widely representative HLA class II alleles DRB1*03:01, DRB1*07:01, DRB1*15:01, DRB3*01:01, DRB3*02:02, DRB4*01:01 and DRB5*01:01 using IEDB-AR^17^. 126444 T cell epitopes were found in unique variants of MSP2 sequenced for Tanzania isolates (Supplementary Table 5). From these, 2217 peptides (Supplementary Table 6) were non-redundant while 29 epitopes out of the 2217 were predicted as strong binders based on an adjusted rank equal or lower than 1% (99^th^ binding percentile) (Supplementary Table 7).

### An iterative analysis pipeline streamlines the processing and interpretation of epidemiological and antigenic diversity data from demultiplexed FASTQ files

In order to streamline the analysis of *msp2* genotyping data, a fully-automated pipeline was assembled in Galaxy, a language-agnostic platform that integrates tools written in R, Python and other programming languages (https://github.com/dfplazag/CCS-Pfal-msp2.git). We observed that the size variant miscalls with the highest read counts corresponded to sequences up to 3 bases longer or shorter than the expected correct calls (Supplementary Fig. 6a). To reduce the read coverage for these miscalls, an iterative noise reduction strategy was applied where sequences called on a first iteration of the pipeline were used to remove from the original dataset highly similar sequences (up to 3 mismatches based on the 4-mismatch difference for the *msp2*s in the strains CD01 and HB3, Supplementary Fig. 6b) which were also up to 3 nucleotides longer or shorter than the sequence called. This approach resulted in a read count reduction of more than 50% for 13 out of the 22 most prevalent size variant miscalls in the dataset (Supplementary Fig. 6c) and a small increase in the number of correct size variant calls from 348 without the noise reduction module to 360 in the set of 235 mock infections. The pipeline is composed of two workflows. The first workflow (Supplementary Fig. 7) takes a collection of demultiplexed FASTQ files and a table with sample names to deliver a tabular dataset including every read assigned to a sample for all the samples in the library and a table with the number of reads for each sample. Workflow 1 also calculates the baseline read count for a sample to be considered *msp2-*positive based on the mean number of reads miss-assigned to water controls plus 2 SDs. Outputs from workflow 1, in addition to a FASTA file with the *msp2* ortholog in *P. billcollinsi* and a second FASTA with the size variant genotyping oligos for *msp2* designed by us and others are the inputs for a second analysis workflow (Fig. 5a). Workflow 2 (Supplementary Fig. 8) runs in two iterations with the noise reduction module filtering out common size variant miscalls between the first and the second iterations (Fig. 5b). While both iterations call size and sequence variants, only iteration two proceeds to construct nucleotide and protein sequence alignments and phylogenies, compute read coverage matrices for size variants in all the samples genotyped, predict T cell epitopes and calculate MOI values for every sample. Workflow 2 can be easily customized to the study of other polymorphic antigens in *P. falciparum* and other pathogens.

**Fig 5.**
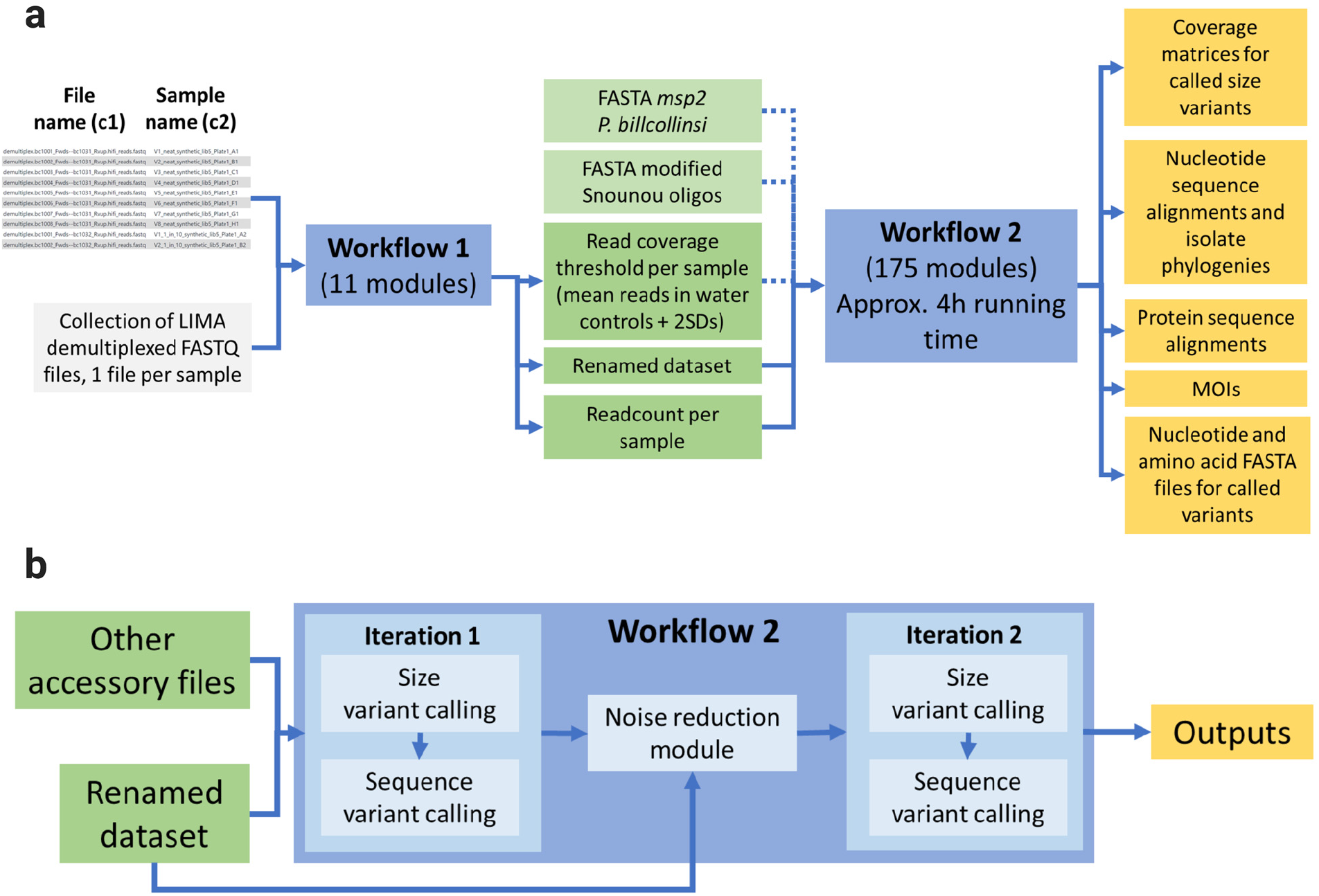
An iterative analysis pipeline streamlines the processing and interpretation of epidemiological and antigenic diversity data from demultiplexed FASTQ files. **a**, General structure of the end-to-end analysis pipeline constructed for *msp2* sequencing data in the bioinformatics analysis interface Galaxy. The pipeline uses as sole inputs a collection of demultiplexed FASTQ files, and three accessory files used for size variant genotyping, file renaming and the construction of isolate phylogenies. Different pipeline outputs are also shown (see also Supplementary Fig. 7 and Supplementary Fig. 8) **b**, A more detailed representation of workflow 2 shows the use of iterative noise reduction for the removal from the dataset of common miscalls resulting from the addition or deletion of up to three bases during PCR amplification or sequencing.

## Discussion

Here, we introduce a long-read amplicon sequencing platform that we believe can be transformative in unlocking the genetic diversity of antigens rich in structural variants. Our platform is also ideal to target and assess the diversity in polymorphic antigens of relevance for the development of immunity to human pathogens for which vaccines are not currently available.

We show that sensitivity and specificity for the CCS genotyping platform can be accurately estimated using complex synthetic mock infections. Most errors introduced during sequencing are amended when individual sequences resulting from each polymerase pass on the circularized template are aligned while producing consensus reads. Moreover, the size and sequence variants called for the synthetic sequences allowed us to define coverage thresholds that can be adjusted to eliminate PCR or sequencing artifacts from the dataset at different FPRs. For single-variant mock infections, we called the correct size and sequence at concentrations of 10 copies/μl in all the 10 variants tested and down to 1 copy/μl in 6 out of the 10.

Multiclonal infections represent a significant challenge in the genotyping of pathogen isolates by short-read sequencing. While a logistically cumbersome approach consists of the sorting and sequencing of individual pathogens present in the isolate^18^, long-read sequencing of complete amplicons circumvents the need for sorting and sequencing of individual clones or the use of algorithms that try to differentiate and assemble short reads derived from a mix of variants^19^. The use of CCS in the determination of infection complexity showed that relaxing the stringency in the calling of variants results in a much better match between the expected and the observed number of co-infecting alleles, although compromising the sequence accuracy of the variants called. Variant calling thresholds can therefore be adjusted depending on the intended application, with higher FPRs (e.g. in average, seven clones can be identified for infections with expected MOI: 10 at an FPR: 0.05) used for epidemiological surveillance and lower FPRs (e.g. in average, one clone can be identified for infections with expected MOI: 10 at an FPR: 0.001) applied to antigen discovery and vaccine development.

Phylogeny reconstruction and the study of multiclonal infections are now made possible by the availability of amplicon sequencing data. Phylogenetic analysis of clones recovered from clinical isolates will be a pivotal tool in the genomic surveillance of infectious disease outbreaks. The ease and low cost of sequencing amplicons for thousands of isolates will also allow the study of population structure and gene flows at a scale that is beyond the current data generation and analysis capacity of whole-genome sequencing approaches^20^. In malaria, the use of highly polymorphic *msp2* sequences as clone-specific “fingerprints” will allow the study of recrudescence and treatment failure with unprecedented resolution joining the battery of markers already in use for sequencing-based follow-up of drug resistance in clinical trials such as *cpp, msp7*^12^, *ama1-D3*^21^, *cpmp*^22^ and *csp*^4^.

The analysis of antigenic diversity will support the development of highly polymorphic antigens as vaccine candidates in two different fronts. First, by identifying fully conserved regions that can then be targeted by the host immune response therefore conferring strain-transcending protection. Second, by informing the production of multivalent cocktails including variable domains for dozens or hundreds of antigens.

The high-throughput analysis of T cell epitopes present in indel-rich antigens can be used to study host-pathogen dynamics with exquisite resolution when coupled to the genotyping of HLA^23, 24^ for the individual harboring specific antigen variants.

A complete survey on the antigenic repertoire of highly polymorphic antigens such as MSP2 could inform the development of a strain-transcending polyvalent antigen cocktails. mRNA as an antigen delivery system has been proven to be effective in eliciting powerful and protective immune responses against SARS-CoV-2^25^. In the future, antigen delivery in the form of codon-optimized mRNAs or alternative delivery platforms for multivalent antigen formulations might be capable of eliciting immunity against dozens or even hundreds of circulating antigen variants in a single vaccine shot imitating, in a way, the protection against clinical disease conferred by repeated exposure to multiple infections.

The implementation of Galaxy for cheap, high-performance computing, facilitates the understanding and sharing of non-sensitive isolate data amongst epidemiologists, wet lab scientists and computational biologists alike. Analysis pipelines and data histories can be easily shared since all the storage and computation takes place in the cloud. Furthermore, the decentralization of storage and computation guarantees the robustness of the data processing and analysis infrastructure. Last but not least, cloud computing infrastructure provides an unparalleled platform to foster international collaborations where analyses can be ran and monitored in real time by collaborators anywhere in the world^26^.

In conclusion, our long-read platform for amplicon sequencing, provides an instrumental tool for the study of complex infections and can be easily adapted to the genotyping of other antigens rich in structural variants. Furthermore, the detailed structural information gained from the sequencing of polymorphic vaccine candidates can now inform the design of polyvalent vaccines capable of transcending the limited protection conferred by formulations targeting individual antigen variants.

## Methods

### Clinical P. falciparum isolates

Venous blood samples were collected in Nyamisati, Pwani Region, Tanzania, in 1999 and 2016 where the population has been followed longitudinally since 1985. At the time of the 1999 survey, parasite prevalence was 64.8% by qPCR^27^. By 2016, prevalence had decreased to 10.2%^28^. Ninety six individuals were included from the 1994 survey: 80 previously shown to be qPCR positive for *P. falciparum* malaria and 16 who tested negative^29^. The 25 isolates included from the 2016 survey were positive by microscopy for *P. falciparum*. This study was approved by the Regional Ethical Committee in Stockholm, Sweden, and the Ethical Review Board of the National Institute for Medical Research in Tanzania.

### Construction of mock infections including synthetic sequences from 10 reference msp2 variants

To assess the specificity and sensitivity of sequence and variant calling, genes encoding *msp2* in the *P. falciparum* reference strains HB3 (LR131339 REGION: 250760..251530, FC27, 292 bp CE variant), CD01 (PfCD01_020011700, FC27, 292 bp CE variant), Dd2 (PfDd2_020009600, FC27, 412 bp CE variant), SN01 (PfSN01_020009800, FC27, 292 bp CE variant), KE01 (PfKE01_020009300, FC27, 373 bp CE variant), SD01 (PfSD01_020012300, IC, 434 bp CE variant), GN01 (PfGN01_020012100, IC, 539 bp CE variant), 7G8 (Pf7G8_020011500, IC, 485 bp CE variant), GB4 (PfGB4_020009800, IC, 533 bp CE variant) and 3D7 (PF3D7_0206800, IC, 473 bp CE variant) were chemically synthesized (GeneScript) and inserted into the pUC57 vector by EcoRV (GAT | ATC) blunt ligation. Mixed mock infections were prepared by diluting 1, 10, 100, 1000, 10000 or 100000 copies/µl of pUC57 with the different *msp2* cassettes in water to simulate MOIs of 1 to 10 (Supplementary Table 1). Each mock infection was assigned to a unique barcode combination.

### DNA extraction and complete amplification of the msp2 open reading frame

For all the clinical isolates, DNA was purified from frozen packed cells in EDTA using QIAamp DNA blood mini kit (Qiagen) following the manufacturer instructions. Concentrations were measured by Nanodrop and the DNA was stored at -80C before use. Taking advantage of the high degree of conservation in the 5’ and 3’ ends of the *msp2* gene, one previously described primer for 5’ annealing (msp2_fw: 5’- ATGAAGGTAATTAAAACATTGTCTATTATA-3’)^13^ that encodes *msp2*’s START codon, as well as a newly designed 3’ primer (msp2_rv2: 5’-TTATATGAATATGGCAAAAGATAAAACAA-3’) that anneals to the end of the *msp2* open reading frame and that encodes *msp2*’s STOP codon were used for a first round of amplification in 15µl reactions. Six units of Phusion Hot Start II DNA Polymerase (Thermo Scientific), 1x Phusion HF Buffer (Thermo Scientific), 0.5μM msp2_fw, 0.5μM msp2_rv2 and 1μl purified DNA were mixed in individual wells of 96-well PCR plates. The first amplification program included 5 steps: 1) 98°C for 30s, 2) 98°C for 10s, 3) 58.1°C for 30s, 4) 72°C for 40s and 5) 72°C for 5min, with 40 cycles of steps 2- 4.

### Asymmetrical sample barcoding and pooling

Low parasitemias of *P. falciparum* had been previously shown to require nested amplification to specifically detect rare PCR templates^30^. For this, barcoded oligos were designed for a second round of nested amplification. Forty standard PacBio barcodes from the Sequel_RSII_96_barcodes_v1 collection (barcodes bc1001 to bc1020 in the forward primer and barcodes bc1031 to bc1050 in the reverse) were added to the 5’ end of high-performance liquid chromatography (HPLC)-purified primers annealing directly downstream or upstream of msp2_F and msp2_rv2, respectively (Supplementary Table 8). Combinations of these primers allowed the barcoding of 384 individual samples in four 96-well PCR plates (Supplementary Fig. 9) for each CCS library that was processed. A reaction volume of 50μl was used for the nested amplification. Twenty units of Phusion Hot Start II DNA Polymerase (Thermo Scientific), 1x Phusion HF Buffer (Thermo Scientific), 0.5 μM forward and reverse primers and 1μl template (product from the first PCR) were used for amplicon barcoding. A PCR program with 7 steps was used: 1) 98°C for 30s, 2) 98°C for 10s, 3) 50°C for 30s, 4) 72°C for 40s, 5) 98°C for 10s, 6) 72°C for 40s and 7) 72°C for 5 min; with steps 2-4 ran in 5 cycles (annealing of the 3’ region of the barcoding primer only) followed by 35 cycles of steps 5 and 6 (annealing along the entire length of the barcoding primer). After barcoding, selected wells containing synthetic mock infection and negative controls, were ran on an agarose gel to verify the presence or the lack thereof of PCR products. The 12 wells from individual plate rows were pooled and cleaned (QIAquick PCR Purification Kit, QIAGEN), and the 30μl elutions resulting from every prep were combined and vortexed thoroughly. Fragment sizes for clean-pooled samples were verified by agarose gel electrophoresis. Concentration, as well as 260/280 and 260/230 nm ratios, were measured in a Thermo Scientific Multiskan GO.

### Single Molecule Real Time (SMRT) library construction for CCS and Lima demultiplexing

PacBio libraries were produced using the SMRTbell™ Template Prep Kit 1.0 according to manufacturer’s instructions. In brief, 500 ng of DNA (amplicon) was purified using PB AMPure beads before DNA damage repair and end-repair followed by ligation of hair-pin adaptors to generate a SMRTbell™ library for circular consensus sequencing. The library was then subjected to exo treatment and PB AMPure bead wash for clean-up. Each 384-plex library was sequenced in a single SMRTcell™ 1M for the PacBio Sequel I instrument using a Sequel 3.0 polymerase and 480 minute movie time, allowing for approx. 32 passes of the polymerase on the barcoded *msp2* products (16 passes on the positive strand + 16 passes on the negative strand). CCS reads were assigned to individual samples and primers were computationally trimmed using the Lima package (in asymmetric mode) that is included in the SMRT Link toolbox (version 10.0.0.108728). Isolates collected in Nyamisati were sequenced twice to assess the reproducibility in the calling of size and sequence variants. FASTQ files were deposited in the European Nucleotide Archive^31^ (Study PRJEB46950, samples ERS12142522 to ERS12143174).

### In silico PCR for size variant genotyping

To assess the presence of size variants for the FC27 and IC1 families of *msp2* that could be compared to previous size variant genotyping surveys using nested PCR and CE-based fragment sizing^14^, FASTQ files for individual samples were collapsed into an individual tabular file with the unique read identifier in column 1 and the read sequence in column 2. Positive or negative strand reads were identified and filtered into two separate tables based on the presence of the tags 5’-AATTTCTTTATTTTTGTTACC-*read*- GTTTTAATTTCAGCAACAC-3’ or 5’-GTGTTGCTGAAATTAAAAC-*read*-GGTAACAAAAATAAAGAAATT-3’, respectively. These tags correspond to the annealing regions on the *msp2* sequence for the barcoding oligos used during library construction on the positive or the negative strands of the gene. The two tables were converted to FASTA and negative strand reads were reverse-complemented and concatenated tail-to-head to the reads from the positive strand into a single FASTA file. A blastn-short search optimized for sequences shorter than 50 bases was ran on this FASTA file^32, 33^, using the sequence from the eight oligos widely used for size variant genotyping of *msp2*^13^ and the ten new primers designed for this study as queries. The primers designed for this study incorporate base substitutions observed for the annealing sites in the *msp2* variants of reference strains ML01, GN01, IT, SD01, TG01, GB4, KE01, Dd2, CD01 and HB3 (Supplementary Table 3). Expectation value cutoff was set to 0.001. Since *msp2* is a low complexity locus^34^, low complexity regions were not filtered out for the analysis. Maximum number of hits to include in the output dataset was set to 1’000’000. 95% was used as identity cutoff and gaps were not allowed. A minimum query coverage per alignment was adjusted to 100%. All the alignments meeting the expected value criteria (-max_hsps) were included in the final output. Output format (-outfmt) was set to Tabular (first 12 columns in Supplementary Table 9). This file was split into 18 separate subfiles, each containing the mapping results for individual oligos based on the Query accession column using the *Filter Tabular* function (version 2.0.0.) from Galaxy, filtering by regex expression matching and including lines matching the name of the 18 genotyping oligos (Supplementary Table 3). The results for forward and reverse oligos were inner joined (Both 1^st^ & 2nd file) with the *Join two files* wrapper using the saccver column (unique CCS read identifiers) as the key. Annealing strand for both forward and reverse oligos was calculated following equation (1):

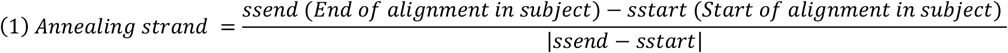

A stringent CCS read filtering process was applied with the following criteria: 1) Number of alignment mismatches (mismatch) = 0 for both forward and reverse oligos, 2) qend = 30 for both forward and reverse oligos, 3) No alignment gaps allowed (gapopen = 0) and 4) Forward and reverse oligos map to opposite strands of the CCS read reflected as a positive strand value for the forward oligo and a negative strand value for the reverse oligo. Size variant calling was computed for each filtered CCS read using equation (2):

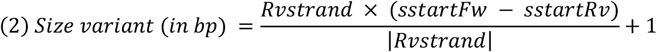

Where:

sstartFw = Start of alignment in subject (CCS read) for the forward primer (ICF or FC27F)

sstartRv = Start of alignment in subject (CCS read) for the reverse primer (ICR or FC27R)

Rvstrand = Annealing strand of the Rv oligo (ICR or FC27R)

Collections of called size variants and their supporting read coverages (proportion of reads in the sample) for all the samples genotyped were collapsed into read coverage matrices using the *Column Join On Multiple Datasets* function on Galaxy. Samples and their called size variants were hierarchically clustered (complete linkage) using Gene Cluster 3.0 with Euclidean as the distance metric. Heat maps were visualized using TreeView 3^35^.

### Size variant genotyping of msp2 by nested PCR and CE

In order to validate size variant calls from *in silico* PCRs on CCS reads, we performed size variant genotyping of *msp2* by nested PCR and fragment sizing by CE as previously described^14^ in a selection of 24 *P. falciparum*-positive isolates. In brief, 20µl reactions to amplify the *msp2* locus were prepared by mixing 0.02 units/μL of AmpliTaq DNA polymerase (ThermoFisher Scientific), 1x AmpliTaq Buffer II (ThermoFisher Scientific), 250nM msp2_fw, 250nM msp2_rv, 2.0mM MgCl2, 125µM dNTP mix and 3μl purified DNA in individual wells of 96-well PCR plates. This first amplification program included 6 steps: 1) 95°C for 5 min, 2) 58°C for 2 min, 3) 72°C for 2 min, 4) 94°C for 1 min, 5) 58°C for 2 min and 6) 72°C for 5 min with 25 cycles of steps 2-4. Next, nested PCR was used to separately amplify VIC- or 6-Carboxyfluorescein-labeled IC1- or FC27-specific products, respectively, in 20µl reactions. These included 0.05 (IC1 reaction) or 0.02 (FC27 reaction) units/μL of AmpliTaq DNA polymerase (ThermoFisher Scientific), 1x AmpliTaq^®^ Buffer II (ThermoFisher Scientific), 300 (IC1 reaction) or 125 (FC27 reaction) nM of forward and reverse 5’-labeled primers (Snounou G *et al*, 1999 in Supplementary Table 3), 1mM MgCl2, 125µM dNTP mix and 1 µl PCR product from the first reaction. The nested amplification program included 6 steps: 1) 95°C for 5 min, 2) 58°C for 1 min, 3) 72°C for 1 min, 4) 94°C for 30s, 5) 58°C for 1 min and 6) 72°C for 5 min with 23 cycles of steps 2-4. Products lengths from the nested reaction were then analyzed in an ABI 3730 PRISM^®^ DNA analyzer. Size variants were called using the GeneMapper® Software 5 and a threshold of 300 relative fluorescence units (RFU).

### Phylogenetic and sequence conservation analysis of isolates in the library pool

All sample-labelled barcode-free reads from clinical isolates were concatenated to the sequence of msp2_fw and the reverse-complemented sequence of msp2_rv2 in the 5’ and 3’ ends, respectively. A FASTA file was created for unique combinations of samples and non-redundant (in the same sample) nucleotide sequences passing the coverage cutoffs for size and sequence variant calling. Sequences were aligned with MAFFT^36^ using the E-INS-i accuracy-oriented method that is suitable for sequences with large unalignable regions (Gap extend penalty 0.0, gap opening penalty 1.53) and sequences in the resulting alignment were reordered according to pairwise similarity. The reference sequence for the ortholog of *msp2* present in *P. billcollinsi* (PlasmoDB: PBILCG01_0209400)^15, 37^ was added to the alignment and used as the tree root in the phylogenetic analysis that followed. The ortholog of *msp2* in *P. billcollinsi* is the only one in the subgenus *Laverania* that is closely related to the *P. falciparum* clade where no subfamily (IC1 or FC27) can be identified from the presence of annealing sites for size variant genotyping oligos (Supplementary Fig. 10). Maximum likelihood trees were constructed with IQ-TREE^38^ using 1000 bootstrap replicates (-bb). 1000 replicates were used for the SH-like approximate likelihood ratio test (SH-aLRT, - alrt) that was calculated for individual tree branches. Trees were visualized and annotated using iTOL v6^39^. Branches supported by more than 80% of the bootstrap replicates were highlighted in the resulting trees. In order to study protein sequence conservation, non-redundant reads in the library passing the variant calling cutoffs were translated using Galaxy *transeq* version 5.0.0 in frame one before computing IC- and FC27-specific alignments with MAFFT (E-INS-I method, gap extend penalty for group-to-group alignment: 0.123, gap opening penalty at group-to-group alignment: 1.53) using a BLOSUM62 matrix. Alignments were visualized with Jalview^40^. Valdar^41^ conservation scores for every residue in the resulting alignments were calculated with AACon^42^. The linear epitopes of well-characterized mouse monoclonal antibodies against the 3D7 and FC27 variants of MSP2^16^ were mapped to the alignment consensus sequences for the IC1 and FC27 families.

### Prediction of T cell epitopes on MSP2 variants in the Immune Epitope Database and Analysis Resource (IEDB-AR)

To test the usefulness of the obtained protein sequences in the study of T cell responses specific to MSP2 variants, HLA class II binding predictions were computed with the MHC-II Binding prediction tool from IEDB-AR^17^ for unique combinations of samples and protein sequences using a seven HLA allele set described to capture 50% of the immune response (DRB1*03:01, DRB1*07:01, DRB1*15:01, DRB3*01:01, DRB3*02:02, DRB4*01:01, DRB5*01:01)^43^. This set has been validated previously in the identification of dominant epitopes independent of ethnicity or HLA variability^44^. Peptide lengths were set to 15 amino acids and epitopes were identified using the consensus method combining four prediction algorithms (NN- align^45^, SMM-align^46^, CombLib and Sturniolo^47^) with top performances^48^. Epitopes with an adjusted rank equal or lower to 1% (99th percentile of binders) were defined as high-affinity binders^44^.

### Construction of an end-to-end pipeline for the analysis of msp2 diversity, epidemiology and antigenicity

To accelerate data processing, variant calling and the generation of downstream epidemiological and antigen biochemistry insights, we designed a fully automated Galaxy-based analysis pipeline made of two separate workflows that run on tandem (Fig. 5). The first workflow takes as inputs a collection of demultiplexed FASTQ files (one file per sample in the sequencing library) and a renaming two-column tabular file with the name of each FASTQ file in the collection in the first column and the names of the matching samples in the second. This first workflow renames the FASTQ files of the sequencing library, counts the number of reads per sample and calculates the minimum number of reads required for a sample to be considered *msp2*-positive based on the average number of reads assigned to the different water controls in the library plus 2 standard deviations. A downstream workflow two takes as inputs a tabular file with the collapsed renamed collection (output from workflow 1), a tabular file with the number of reads per sample (output from workflow 1), a FASTA file with the ortholog gene of *msp2* present in *P. billcollinsi*^37^ (to be used as tree root in the construction of isolate phylogenies) and a FASTA file with the sequences of the oligos used for size variant genotyping. This second workflow filters out *msp2*-negative samples, classifies reads as belonging to the IC1 or FC27 families of *msp2* and calculates size variants for each individual read. A suit of epidemiologically relevant outputs are provided by the pipeline including sequence- and size variant-based number of co-infecting clones (MOI) per sample and maximum likelihood phylogenetic trees for the *msp2* variants present in the library. Finally, the pipeline completes the *msp2* sequence for each read in the dataset by adding to the 5’ and 3’ ends the corresponding oligos used in the first PCR of the library construction protocol and translates the reads into protein sequences that are then used to predict T cell epitopes, construct multiple sequence alignments and calculate linear epitope abundance matrices. Nine versions of workflow two are available using all the variant calling cutoff combinations from Supplementary Table 2 in order to match FPRs for size and sequence variant calling of 0.001, 0.01 and 0.05. All pipelines and accessory files can be found in GitHub (https://github.com/dfplazag/CCS-Pfal-msp2.git).

## Acknowledgements

The authors would like to thank the study participants and research teams contributing to the clinical malaria isolates collected in Tanzania. The project was funded by the Swedish Research Council (VR MH 2018-04468 and 2018-02668) and the Stockholm County Council (Project grant SLL20150135 and FoUI- 953118). The authors would also like to acknowledge the National Genomics Infrastructure (NGI) / Uppsala Genome Center and UPPMAX for providing assistance in sequencing and computational infrastructure. Work performed at NGI / Uppsala Genome Center has been funded by RFI / VR and Science for Life Laboratory, Sweden. The data handling and storage were enabled by resources provided by the Swedish National Infrastructure for Computing (SNIC) at the Uppsala Multidisciplinary Center for Advanced Computational Science partially funded by the Swedish Research Council (VR 2018-05973). The Galaxy server that was used for some calculations is in part funded by Collaborative Research Centre 992 Medical Epigenetics (DFG grant SFB 992/1 2012) and German Federal Ministry of Education and Research (BMBF grants 031 A538A/A538C RBC, 031L0101B/031L0101C de.NBI-epi, 031L0106 de.STAIR (de.NBI)). Finally, we are grateful to Dr. Carlos Fernando Suárez for the insightful discussions on the population dynamics of circulating T cell epitopes from *P. falciparum*.

