## Supplementary_File_1 for "A genomic platform for epidemiological surveillance and vaccine antigen discovery using long-read amplicon sequencing"

**Supplementary information**

**Supplementary Table 2**. Coverage thresholds for size and sequence variant calling at false positive rates of 0.001, 0.01 and 0.05. Since sequence variants are called on reads matching previously called size variants, thresholds for sequence variant calling at specific FPRs vary depending on the thresholds previously used for size variant calling. False negative rates for the same threshold combinations are shown.

| Size variant calling Iteration 2 | | | Sequence variant calling Iteration 2 | | |
|---|---|---|---|---|---|
| **FPR** | **Threshold (% reads in sample)** | **FNR** | **FPR** | **Threshold (% reads in sample)** | **FNR** |
| | | | **0.001** | 16 | 0.739 |
| **0.001** | 14 | 0.688 | **0.01** | 6.6 | 0.679 |
| | | | **0.05** | 2.4 | 0.658 |
| | | | **0.001** | 12 | 0.734 |
| **0.01** | 3.8 | 0.479 | **0.01** | 5.3 | 0.540 |
| | | | **0.05** | 1.8 | 0.484 |
| | | | **0.001** | 9.6 | 0.711 |
| **0.05** | 1.5 | 0.362 | **0.01** | 4.4 | 0.584 |
| | | | **0.05** | 1.6 | 0.444 |

**Supplementary Table 3**. Primers for size variant genotyping of the IC1 and FC27 families of *msp2*. Genotyping by capillary electrophoresis was carried out using the original oligos from Snounou G *et al*, 1999. In addition, new forward primers were designed based on the more recent availability of reference *msp2* sequences ML01, GN01, IT, SD01, TG01, GB4, KE01, Dd2, CD01, SN01 and HB3. These newly designed primers were only used for *in silico* PCR-based size variant genotyping.

| Primer name | Primer sequence 5' → 3' |
|---|---|
| ICF1 (Snounou G *et al*, 1999) | AGAAGTATGGCAGAAAGTAAGCCTCCTACT |
| ICF2 (Snounou G *et al*, 1999) | AGAAGTATGGCAGAAAGTAATCCTCCTACT |
| ICF3 (Snounou G *et al*, 1999) | AGAAGTATGGCAGAAAGTAAGCCTTCTACT |
| ICF4 (Snounou G *et al*, 1999) | AGAAGTATGGCAGAAAGTAATCCTTCTACT |
| ICF_ML01 (Designed for this study) | AGAAGTATGGAAGAAAGTAATCCTCCTACT |
| ICF_GN01 (Designed for this study) | AGAAGTATGGCAGTAAGTAATCCTTCTACT |
| ICF_IT (Designed for this study) | AGAAGTATGACAGAAAGTAATCCTCCTACT |
| ICF_SD01 (Designed for this study) | AGAAGTATGTCAGAAAGTAAGCCTCCTACT |
| ICF_TG01 (Designed for this study) | AGAAGTATGACAGAAAGTAAGCCTCCTACT |
| ICF_GB4 (Designed for this study) | AGAAGTATGGCAGAAAGTAAGACTCCTACT |
| ICR (Snounou G *et al*, 1999) | GATTGTAATTCGGGGGATTCAGTTTGTTCG |
| FC27F1 (Snounou G *et al*, 1999) | AATACTAAGAGTGTAGGTGCAAATGCTCCA |
| FC27F2 (Snounou G *et al*, 1999) | AATACTAAGAGTGTAGGTGCAGATGCTCCA |
| FC27F_KE01 (Designed for this study) | ACTACTAATAGTGTAGATGCAAATGCTCCA |
| FC27F_Dd2 (Designed for this study) | AATACTACTAGTGTAGGTGCAAATGCTCCA |
| FC27F_CD01_SN01 (Designed for this study) | AATACTAATAGTGTAGGTGCAGATGCTCCA |
| FC27F_HB3 (Designed for this study) | AATACTAAGAGTGTAGGTGCAAATGCTCCA |
| FC27R (Snounou G *et al*, 1999) | TTTTATTTGGTGCATTGCCAGAACTTGAAC |

**Supplementary Table 4**. Size variant calling for *msp2* by CCS and CE in a selection of 24 *msp2*-positive clinical isolates from Tanzania. Size variants are presented in bp. Parentheses show read coverages and RFU values for size variants called by CCS and CE, respectively. Matching size variants for the two methods are highlighted in bold.

| Sample | CCS sizevar calls in bp (%reads in sample) | CE sizevar calls in bp (RFUs) |
|---|---|---|
| 103_Nya_2016_lib6_Plate4_F3 | **IC_503. (100.0)** | **IC_507 (6347)** |
| 104_Nya_2016_lib6_Plate4_G3 | **IC_527. (93.8)** | **IC_525 (23252)** |
| 123_Nya_2016_lib6_Plate4_D4 | **FC27_328. (91.4)** | **FC_329 (12886)**, IC_553 (343) |
| 137_Nya_2016_lib6_Plate4_G4 | FC27_328. (100) | IC_533 (439), IC_543 (390) |
| 141_Nya_2016_lib6_Plate4_H4 | FC27_364. (14.3), IC_470. (71.4), IC_556. (14.3) | FC_298 (588), IC_518 (528), IC_528 (548) |
| 146_Nya_2016_lib6_Plate4_B5 | FC27_292. (50.0), **FC27_328. (16.7)**, IC_473. (16.7), **IC_535. (16.7)** | **FC_329 (14690)**, **IC_542 (1055)** |
| 160_Nya_2016_lib6_Plate4_E5 | **IC_611. (93.9)** | IC_586 (377), **IC_609 (8682)** |
| 163_Nya_2016_lib6_Plate4_F5 | IC_470. (40.0), IC_611. (26.7) | IC_504 (25475) |
| 173_Nya_2016_lib6_Plate4_G5 | **IC_497. (21.6)**, **IC_533. (56.8)** | FC_403 (577), IC_427 (323), **IC_499 (10185)**, **IC_533 (9003)**, IC_544 (12253) |
| 178_Nya_2016_lib6_Plate4_H5 | **IC_587. (96.0)** | FC_412 (16466), IC_379 (1098), IC_403 (1190), IC_456 (1050), **IC_583 (11539)**, IC_604 (5914) |
| 310_Nya_2016_lib6_Plate4_D6 | **IC_680. (40.6)** | FC_293 (4668), FC_329 (5151), FC_413 (6129), FC_484 (371), IC_332 (420), IC_491 (5168), IC_535 (2935), IC_575 (4694), **IC_677 (9689)** |
| 33_Nya_2016_lib6_Plate4_F1 | FC27_301. (84.0) | FC_329 (18454), IC_618 |
| 365_Nya_2016_lib6_Plate4_A7 | FC27_256. (81.2) | IC_473 (13287), IC_544 (1331) |
| 398_Nya_2016_lib6_Plate4_B7 | FC27_256. (50.0), FC27_292. (20.0) | FC_549 (16953), IC_486 (1103) |
| 409_Nya_2016_lib6_Plate4_C7 | **FC27_256. (97.6)** | **FC_255 (11106)**, IC_488 (476), IC_561 (762) |
| 417_Nya_2016_lib6_Plate4_D7 | **IC_539. (95.3)** | FC_249 (763), **IC_538 (11300)** |
| 429_Nya_2016_lib6_Plate4_E7 | FC27_256. (46.4), **FC27_328. (35.7)** | FC_316 (6492), **FC_329 (8882)**, IC_527 (15600) |
| 432_Nya_2016_lib6_Plate4_F7 | **FC27_256. (93.6)** | **FC_255 (15010)**, IC_525 (12809) |
| 461_Nya_2016_lib6_Plate4_G7 | FC27_256. (42.3), FC27_292. (31.0) | FC_240 (680), IC_526 (12790), IC_582 (363) |
| 58_Nya_2016_lib6_Plate4_H1 | FC27_301. (50.0), IC_437. (50.0) | FC_315 (11753), IC_513 (662) |
| 83_Nya_2016_lib6_Plate4_F2 | **IC_527. (89.9)** | **IC_527 (9347)** |
| 86_Nya_2016_lib6_Plate4_H2 | **IC_497. (59.0)** | FC_412 (15004), IC_471 (5845), **IC_490 (6037)**, **IC_499 (12138)** |
| 95_Nya_2016_lib6_Plate4_B3 | IC_464. (100.0) | IC_785 (10415) |
| 97_Nya_2016_lib6_Plate4_D3 | **IC_464. (92.3)** | FC_329 (14464), **IC_465 (10839)**, IC_582 (2286) |

**Supplementary Table 8**. Oligos used for asymmetrical barcoding of *msp2* long amplicons in the nested PCR reaction

| Primer name | PacBio standard Barcode | Annealing sequence to *msp2* | Oligos used for the nested PCR |
|---|---|---|---|
| bc1001_Fwds | CACATATCAGAGTGCG | AATTTCTTTATTTTTGTTACC | CACATATCAGAGTGCGAATTTCTTTATTTTTGTTACC |
| bc1002_Fwds | ACACACAGACTGTGAG | AATTTCTTTATTTTTGTTACC | ACACACAGACTGTGAGAATTTCTTTATTTTTGTTACC |
| bc1003_Fwds | ACACATCTCGTGAGAG | AATTTCTTTATTTTTGTTACC | ACACATCTCGTGAGAGAATTTCTTTATTTTTGTTACC |
| bc1004_Fwds | CACGCACACACGCGCG | AATTTCTTTATTTTTGTTACC | CACGCACACACGCGCGAATTTCTTTATTTTTGTTACC |
| bc1005_Fwds | CACTCGACTCTCGCGT | AATTTCTTTATTTTTGTTACC | CACTCGACTCTCGCGTAATTTCTTTATTTTTGTTACC |
| bc1006_Fwds | CATATATATCAGCTGT | AATTTCTTTATTTTTGTTACC | CATATATATCAGCTGTAATTTCTTTATTTTTGTTACC |
| bc1007_Fwds | TCTGTATCTCTATGTG | AATTTCTTTATTTTTGTTACC | TCTGTATCTCTATGTGAATTTCTTTATTTTTGTTACC |
| bc1008_Fwds | ACAGTCGAGCGCTGCG | AATTTCTTTATTTTTGTTACC | ACAGTCGAGCGCTGCGAATTTCTTTATTTTTGTTACC |
| bc1009_Fwds | ACACACGCGAGACAGA | AATTTCTTTATTTTTGTTACC | ACACACGCGAGACAGAAATTTCTTTATTTTTGTTACC |
| bc1010_Fwds | ACGCGCTATCTCAGAG | AATTTCTTTATTTTTGTTACC | ACGCGCTATCTCAGAGAATTTCTTTATTTTTGTTACC |
| bc1011_Fwds | CTATACGTATATCTAT | AATTTCTTTATTTTTGTTACC | CTATACGTATATCTATAATTTCTTTATTTTTGTTACC |
| bc1012_Fwds | ACACTAGATCGCGTGT | AATTTCTTTATTTTTGTTACC | ACACTAGATCGCGTGTAATTTCTTTATTTTTGTTACC |
| bc1013_Fwds | CTCTCGCATACGCGAG | AATTTCTTTATTTTTGTTACC | CTCTCGCATACGCGAGAATTTCTTTATTTTTGTTACC |
| bc1014_Fwds | CTCACTACGCGCGCGT | AATTTCTTTATTTTTGTTACC | CTCACTACGCGCGCGTAATTTCTTTATTTTTGTTACC |
| bc1015_Fwds | CGCATGACACGTGTGT | AATTTCTTTATTTTTGTTACC | CGCATGACACGTGTGTAATTTCTTTATTTTTGTTACC |
| bc1016_Fwds | CATAGAGAGATAGTAT | AATTTCTTTATTTTTGTTACC | CATAGAGAGATAGTATAATTTCTTTATTTTTGTTACC |
| bc1017_Fwds | CACACGCGCGCTATAT | AATTTCTTTATTTTTGTTACC | CACACGCGCGCTATATAATTTCTTTATTTTTGTTACC |
| bc1018_Fwds | TCACGTGCTCACTGTG | AATTTCTTTATTTTTGTTACC | TCACGTGCTCACTGTGAATTTCTTTATTTTTGTTACC |
| bc1019_Fwds | ACACACTCTATCAGAT | AATTTCTTTATTTTTGTTACC | ACACACTCTATCAGATAATTTCTTTATTTTTGTTACC |
| bc1020_Fwds | CACGACACGACGATGT | AATTTCTTTATTTTTGTTACC | CACGACACGACGATGTAATTTCTTTATTTTTGTTACC |
| bc1031_Rvup | GATGTCTGAGTGTGTG | GTGTTGCTGAAATTAAAAC | GATGTCTGAGTGTGTGGTGTTGCTGAAATTAAAAC |
| bc1032_Rvup | GAGACTAGAGATAGTG | GTGTTGCTGAAATTAAAAC | GAGACTAGAGATAGTGGTGTTGCTGAAATTAAAAC |
| bc1033_Rvup | TCTCGTCGCAGTCTCT | GTGTTGCTGAAATTAAAAC | TCTCGTCGCAGTCTCTGTGTTGCTGAAATTAAAAC |
| bc1034_Rvup | ATGTGTATATAGATAT | GTGTTGCTGAAATTAAAAC | ATGTGTATATAGATATGTGTTGCTGAAATTAAAAC |
| bc1035_Rvup | GCGCGCGCACTCTCTG | GTGTTGCTGAAATTAAAAC | GCGCGCGCACTCTCTGGTGTTGCTGAAATTAAAAC |
| bc1036_Rvup | GAGACACGTCGCACAC | GTGTTGCTGAAATTAAAAC | GAGACACGTCGCACACGTGTTGCTGAAATTAAAAC |
| bc1037_Rvup | ACACATATCGCACTAC | GTGTTGCTGAAATTAAAAC | ACACATATCGCACTACGTGTTGCTGAAATTAAAAC |
| bc1038_Rvup | GTGTGTCTCGATGCGC | GTGTTGCTGAAATTAAAAC | GTGTGTCTCGATGCGCGTGTTGCTGAAATTAAAAC |
| bc1039_Rvup | CGCACACATAGATACA | GTGTTGCTGAAATTAAAAC | CGCACACATAGATACAGTGTTGCTGAAATTAAAAC |
| bc1040_Rvup | TGTCATATGAGAGTGT | GTGTTGCTGAAATTAAAAC | TGTCATATGAGAGTGTGTGTTGCTGAAATTAAAAC |
| bc1041_Rvup | TCTCGCGCGTGCACGC | GTGTTGCTGAAATTAAAAC | TCTCGCGCGTGCACGCGTGTTGCTGAAATTAAAAC |
| bc1042_Rvup | CTCGCTCGACGAGCGC | GTGTTGCTGAAATTAAAAC | CTCGCTCGACGAGCGCGTGTTGCTGAAATTAAAAC |
| bc1043_Rvup | TATAGAGCTCTACATA | GTGTTGCTGAAATTAAAAC | TATAGAGCTCTACATAGTGTTGCTGAAATTAAAAC |
| bc1044_Rvup | GCTGAGACGACGCGCG | GTGTTGCTGAAATTAAAAC | GCTGAGACGACGCGCGGTGTTGCTGAAATTAAAAC |
| bc1045_Rvup | ACATATCGTACTCTCT | GTGTTGCTGAAATTAAAAC | ACATATCGTACTCTCTGTGTTGCTGAAATTAAAAC |
| bc1046_Rvup | GATATATCGAGTATAT | GTGTTGCTGAAATTAAAAC | GATATATCGAGTATATGTGTTGCTGAAATTAAAAC |
| bc1047_Rvup | TGTCATGTGTACACAC | GTGTTGCTGAAATTAAAAC | TGTCATGTGTACACACGTGTTGCTGAAATTAAAAC |
| bc1048_Rvup | GTGTGCACTCACACTC | GTGTTGCTGAAATTAAAAC | GTGTGCACTCACACTCGTGTTGCTGAAATTAAAAC |
| bc1049_Rvup | ACACGTGTGCTCTCTC | GTGTTGCTGAAATTAAAAC | ACACGTGTGCTCTCTCGTGTTGCTGAAATTAAAAC |
| bc1050_Rvup | GATATACGCGAGAGAG | GTGTTGCTGAAATTAAAAC | GATATACGCGAGAGAGGTGTTGCTGAAATTAAAAC |

**Supplementary Table 9**. Blastn data retrieved for the size variant genotyping of CCS reads (columns 1 to 12) and the noise reduction module of the analysis pipeline (columns 1 to 25)

| Column | NCBI name | Description |
|---|---|---|
| 1 | qaccver | Query accession dot version (name of the genotyping oligo) |
| 2 | saccver | Subject accession dot version (CCS database read hit) |
| 3 | pident | Percentage of identical matches |
| 4 | length | Alignment length |
| 5 | mismatch | Number of mismatches |
| 6 | gapopen | Number of gap openings |
| 7 | qstart | Start of alignment in query (genotyping oligo) |
| 8 | qend | End of alignment in query (genotyping oligo) |
| 9 | sstart | Start of alignment in subject (CCS read) |
| 10 | send | End of alignment in subject (CCS read) |
| 11 | evalue | Expectation value (E-value) |
| 12 | bitscore | Bit score |
| 13 | sallseqid | All subject Seq-id(s), separated by a ';' |
| 14 | score | Raw score |
| 15 | nident | Number of identical matches |
| 16 | positive | Number of positive-scoring matches |
| 17 | gaps | Total number of gaps |
| 18 | ppos | Percentage of positive-scoring matches |
| 19 | qframe | Query frame |
| 20 | sframe | Subject frame |
| 21 | qseq | Aligned part of query sequence |
| 22 | sseq | Aligned part of subject sequence |
| 23 | qlen | Query sequence length |
| 24 | slen | Subject sequence length |
| 25 | salltitles | All subject title(s), separated by a '<>' |

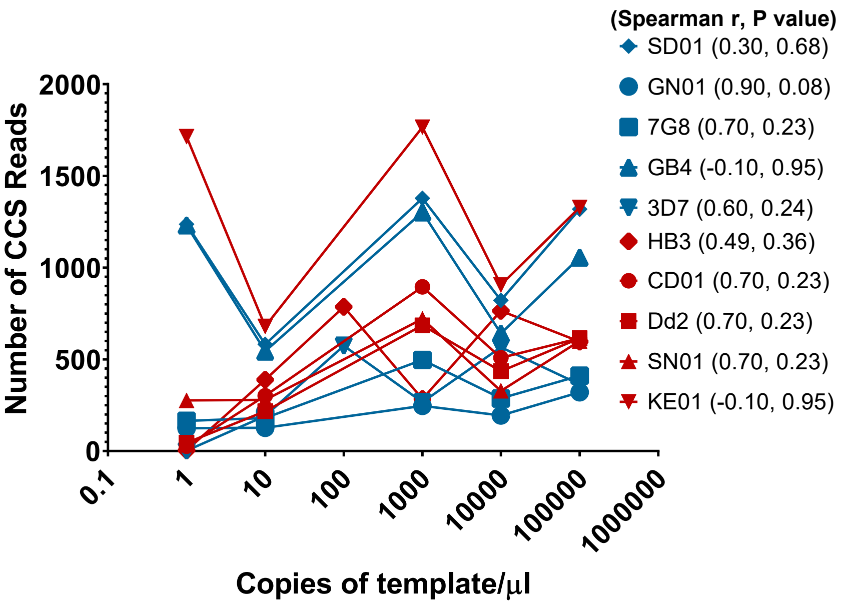


**Supplementary Fig. 1. Lack of correlation between template concentration and read counts per sample**. Number of reads per sample in mock infections including single clones at concentrations ranging from 1 to 100000 copies/µl are presented. Spearman R and P values per *msp2* variant are shown in parentheses. Variants of the IC and FC27 families are shown in blue and red, respectively

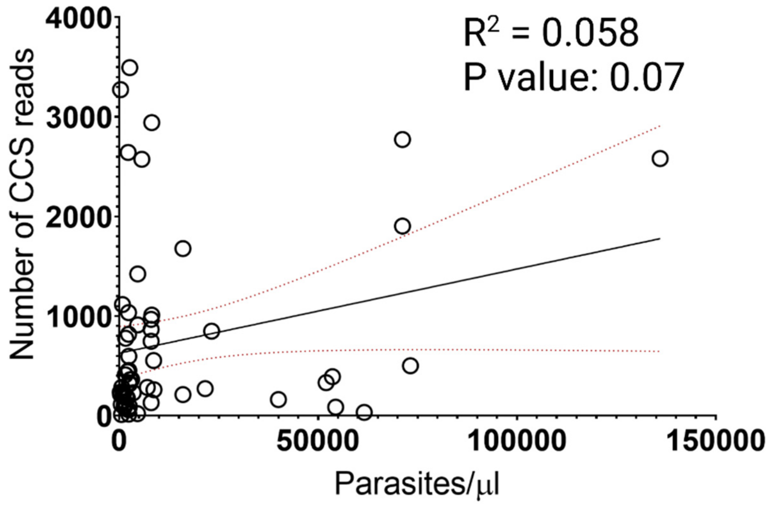


**Supplementary Fig. 2. Parasite density and read counts for 58 microscopy-positive isolates from Tanzania**. There is no correlation between parasite density in the isolate as measured by microscopy and read coverage per sample. Spearman r: 0.2568. R square, trend line (solid) for a simple linear regression and 95% confidence intervals (doted lines) are shown.

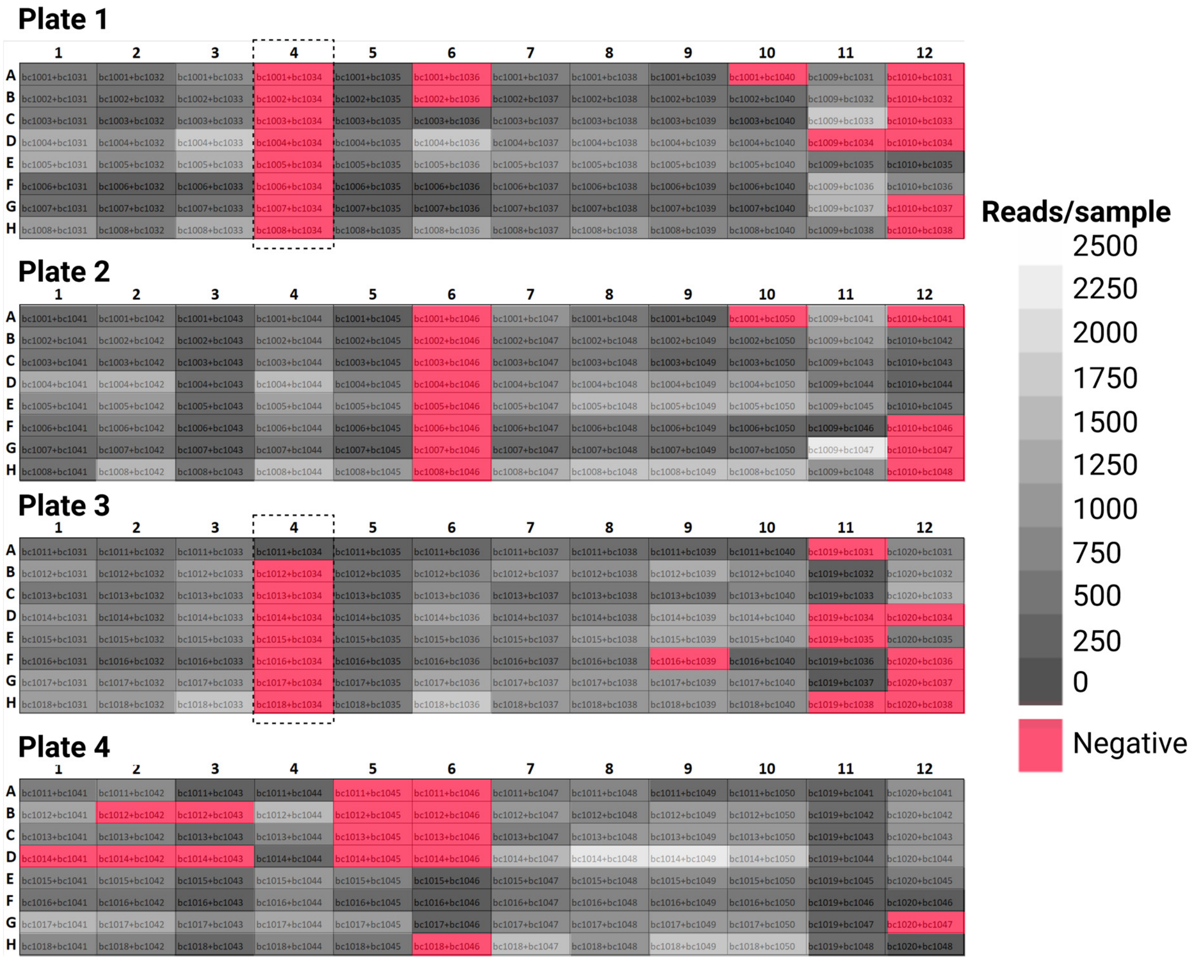


Library 5, reads/sample for samples with more than or equal to 124 reads.

**Supplementary Fig. 3. Reads per sample in the four plates of the synthetic mock infection library.** Wells were classified as negative if they had less than the 124 reads cutoff established from the water controls. 324 samples in the library (84.4% of all the samples) yielded read counts above the background cutoff. Samples barcoded with bc1034 in plates 1 and 3 are shown within dashed frames. bc1034 was used in the barcoding of the 100 copies/µl dilution for single-variant mock infections of the reference *msp2*s SD01, GN01, 7G8, GB4, CD01, Dd2, SN01 and KE01.

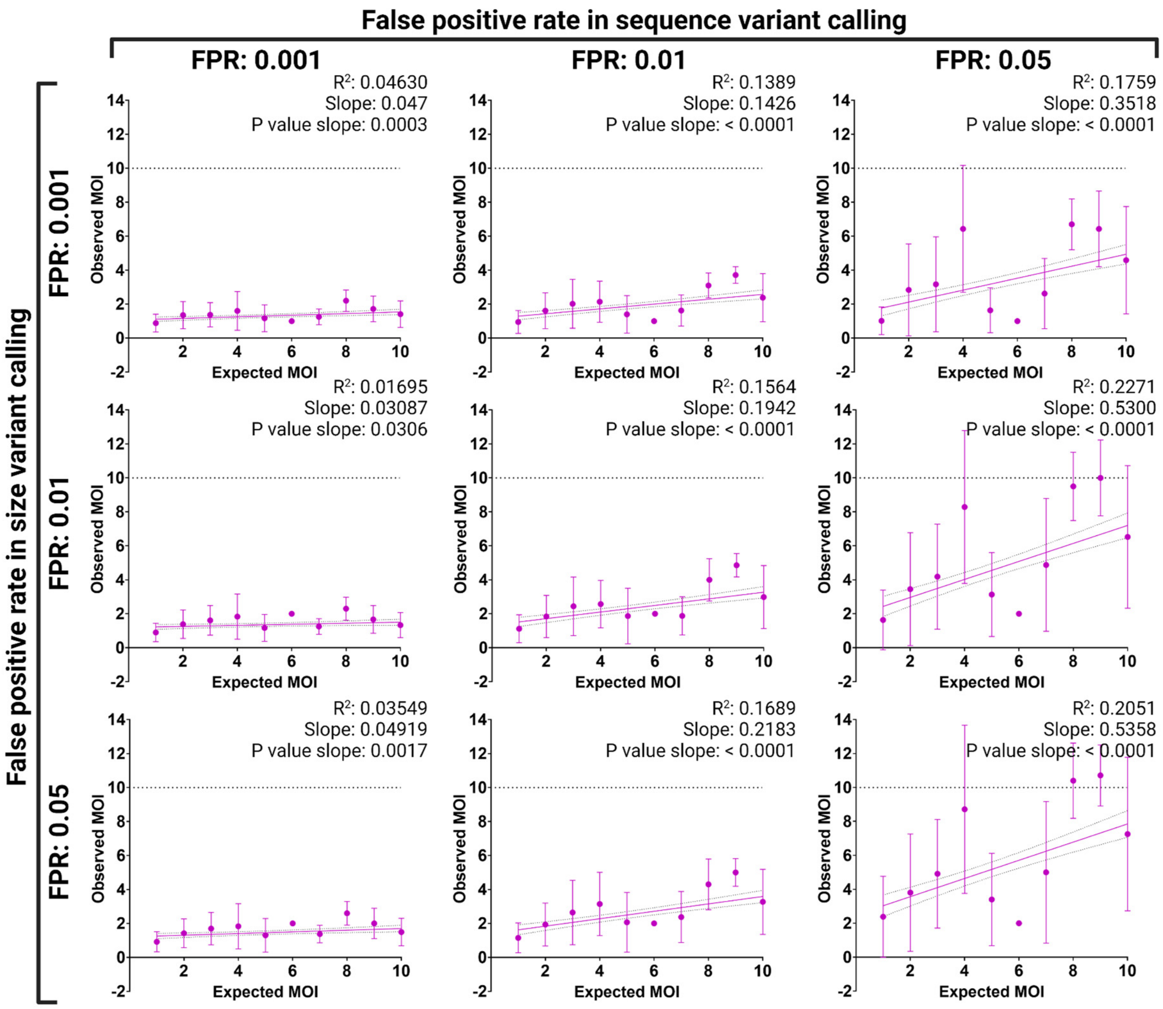


**Supplementary Fig. 4. Observed multiplicity of infection as a function of the FPRs at different variant calling thresholds**. The selection of variant calling thresholds resulting in different FPRs affect the amount of clones that can be detected per isolate. R squared, slope, P value and 95% confidence intervals for simple linear regressions fitted to the 9 possible FRP scenarios are included. Mean and standard deviations (error bars) are also shown. A dashed line marks the maximum number of observed variants possible (MOI: 10) for the prepared mock infections.

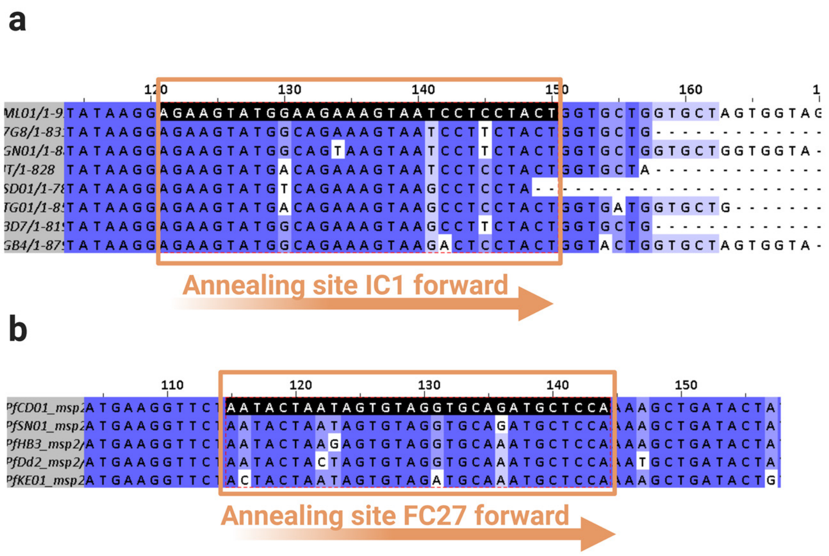


**Supplementary Fig. 5. Annealing site for size variant genotyping primers.** (forward only, no new substitutions were observed in reverse primers) on **a,** IC1 and **b,** FC27 *msp2* variants sequenced after 1999 when the first genotyping oligos for the gene were designed based on the variants 3D7 and FC27 only (Snounou G *et al*, 1999).

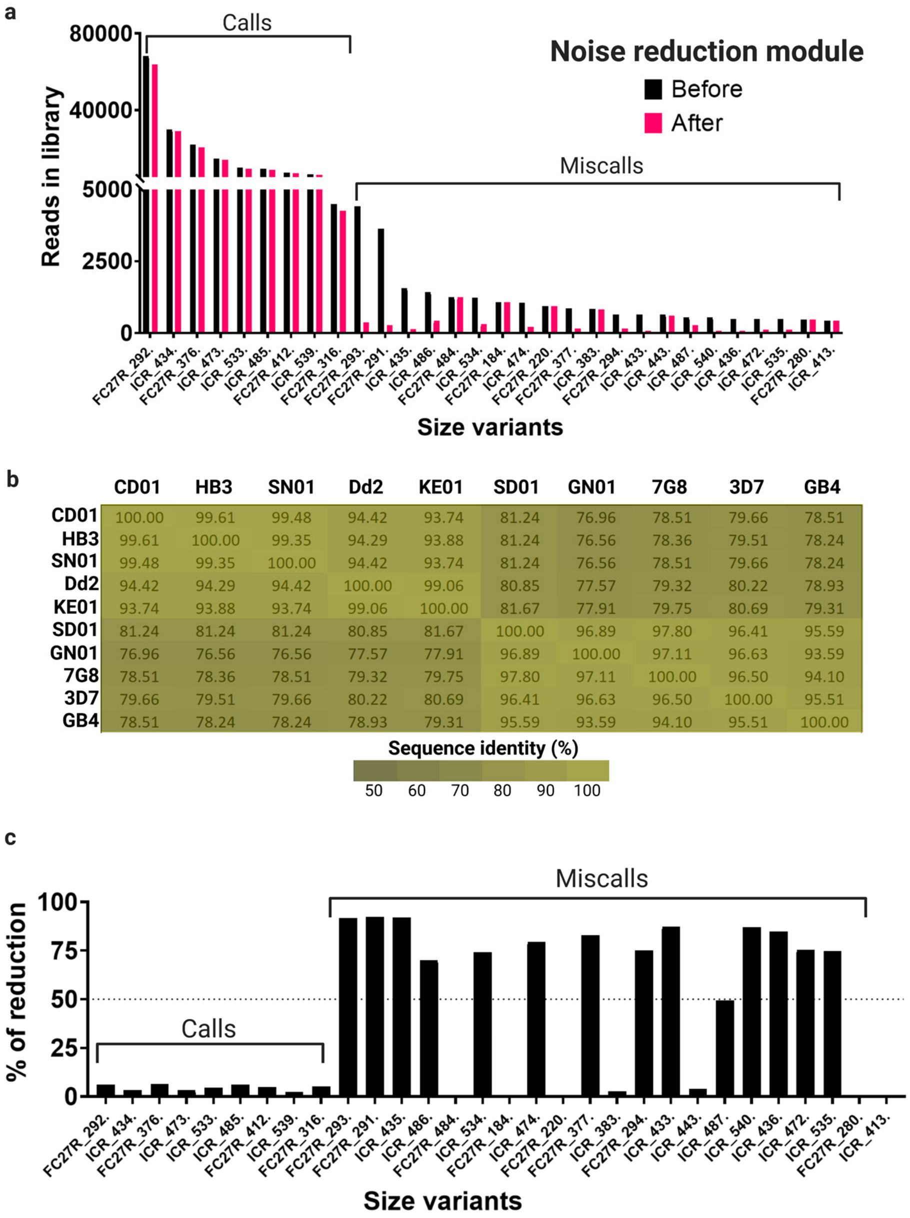


**Supplementary Fig. 6. Quantification of iterative noise reduction for miscalls with the addition or deletion of up to 3 nucleotides resulting from PCR or sequencing errors**. **a,** Number of reads for the 30 most common size variants in the synthetic mock infection library before (black) and after (fuchsia) noise reduction. Correct calls show the highest read counts in the library. **b,** Sequence identity matrix for the same 10 *msp2* variants. **c,** Reduction percentage in read counts for the 30 most common size variants in the same library. A reduction of more than 50% in the assigned number of reads can be observed as a result of the noise reduction module for 13 out of the 22 most prevalent miscalls in the dataset.

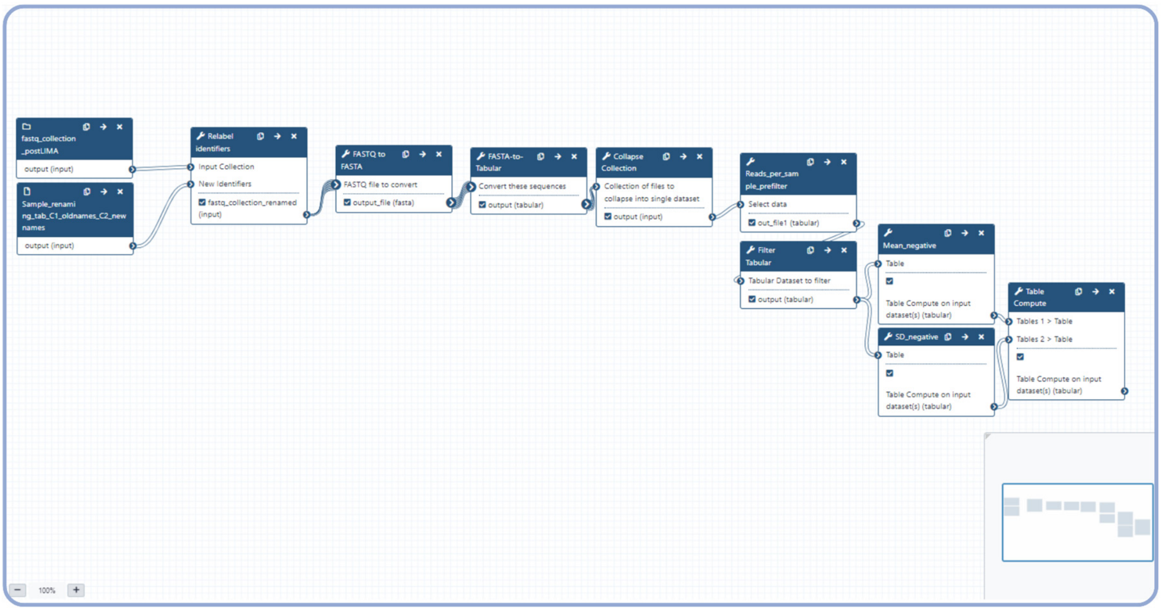


**Supplementary Fig. 7. Galaxy workflow 1.** This workflow produces a tabular dataset collapsed to include every read assigned to a sample for all the samples in the library and a table with the number of reads for each sample. Workflow 1 also calculates the baseline read count for a sample to be considered *msp2*-positive based on the mean number of reads miss-assigned to water controls plus 2 SDs (https://github.com/dfplazag/CCS-Pfal-msp2.git).

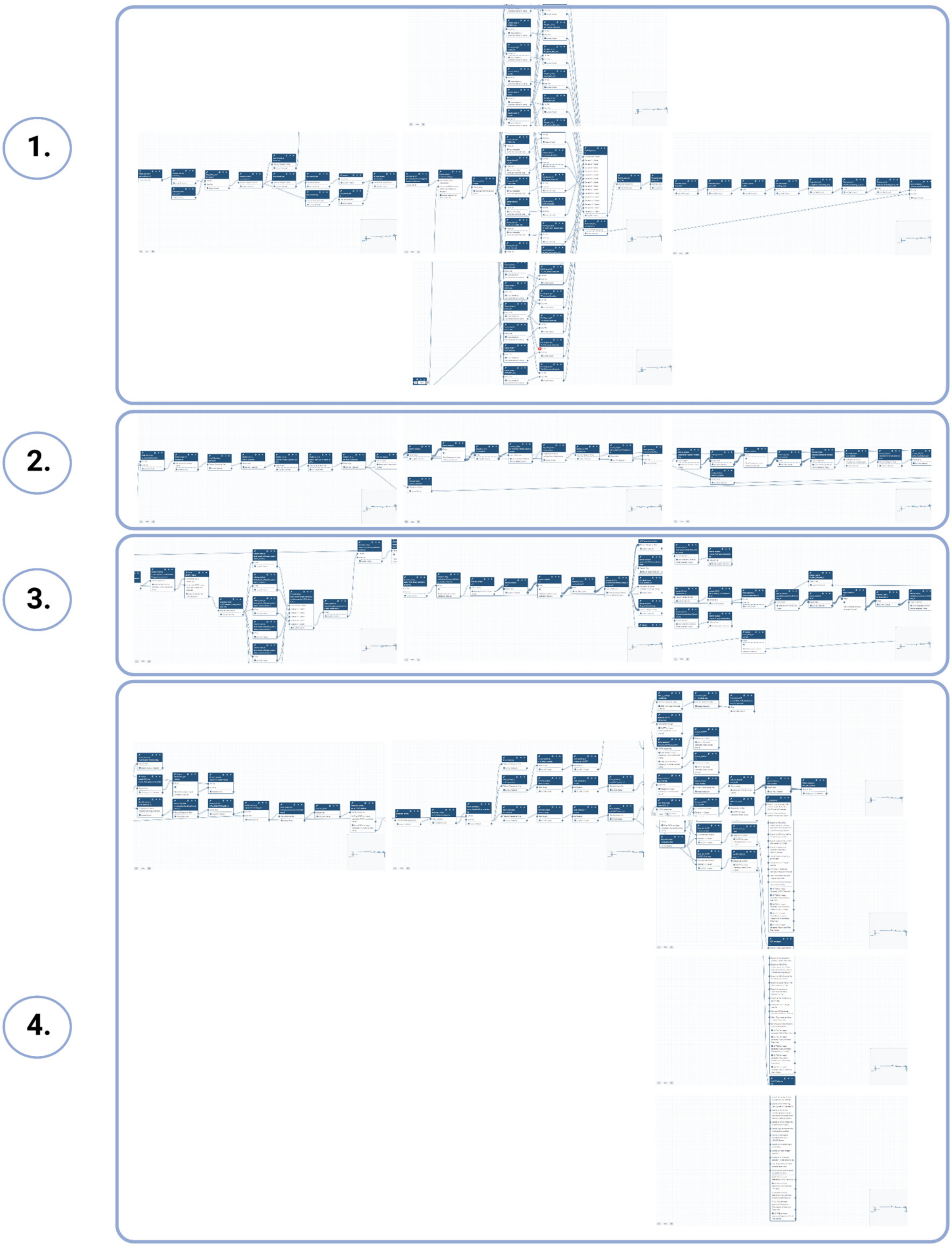


**Supplementary Fig. 8. Galaxy workflow 2.** Workflow 2 takes two accessory files and the outputs from workflow 1 to construct nucleotide and protein sequence alignments and phylogenies, compute read coverage matrices for size variants in all the samples genotyped, predict T cell epitopes and calculate MOI values for every sample (https://github.com/dfplazag/CCS-Pfal-msp2.git).

**Plate 1**

| | 1 | 2 | 3 | 4 | 5 | 6 | 7 | 8 | 9 | 10 | 11 | 12 |
|---|---|---|---|---|---|---|---|---|---|---|---|---|
| A | bc1001+bc1031 | bc1001+bc1032 | bc1001+bc1033 | bc1001+bc1034 | bc1001+bc1035 | bc1001+bc1036 | bc1001+bc1037 | bc1001+bc1038 | bc1001+bc1039 | bc1001+bc1040 | bc1009+bc1031 | bc1010+bc1031 |
| B | bc1002+bc1031 | bc1002+bc1032 | bc1002+bc1033 | bc1002+bc1034 | bc1002+bc1035 | bc1002+bc1036 | bc1002+bc1037 | bc1002+bc1038 | bc1002+bc1039 | bc1002+bc1040 | bc1009+bc1032 | bc1010+bc1032 |
| C | bc1003+bc1031 | bc1003+bc1032 | bc1003+bc1033 | bc1003+bc1034 | bc1003+bc1035 | bc1003+bc1036 | bc1003+bc1037 | bc1003+bc1038 | bc1003+bc1039 | bc1003+bc1040 | bc1009+bc1033 | bc1010+bc1033 |
| D | bc1004+bc1031 | bc1004+bc1032 | bc1004+bc1033 | bc1004+bc1034 | bc1004+bc1035 | bc1004+bc1036 | bc1004+bc1037 | bc1004+bc1038 | bc1004+bc1039 | bc1004+bc1040 | bc1009+bc1034 | bc1010+bc1034 |
| E | bc1005+bc1031 | bc1005+bc1032 | bc1005+bc1033 | bc1005+bc1034 | bc1005+bc1035 | bc1005+bc1036 | bc1005+bc1037 | bc1005+bc1038 | bc1005+bc1039 | bc1005+bc1040 | bc1009+bc1035 | bc1010+bc1035 |
| F | bc1006+bc1031 | bc1006+bc1032 | bc1006+bc1033 | bc1006+bc1034 | bc1006+bc1035 | bc1006+bc1036 | bc1006+bc1037 | bc1006+bc1038 | bc1006+bc1039 | bc1006+bc1040 | bc1009+bc1036 | bc1010+bc1036 |
| G | bc1007+bc1031 | bc1007+bc1032 | bc1007+bc1033 | bc1007+bc1034 | bc1007+bc1035 | bc1007+bc1036 | bc1007+bc1037 | bc1007+bc1038 | bc1007+bc1039 | bc1007+bc1040 | bc1009+bc1037 | bc1010+bc1037 |
| H | bc1008+bc1031 | bc1008+bc1032 | bc1008+bc1033 | bc1008+bc1034 | bc1008+bc1035 | bc1008+bc1036 | bc1008+bc1037 | bc1008+bc1038 | bc1008+bc1039 | bc1008+bc1040 | bc1009+bc1038 | bc1010+bc1038 |

**Plate 2**

| | 1 | 2 | 3 | 4 | 5 | 6 | 7 | 8 | 9 | 10 | 11 | 12 |
|---|---|---|---|---|---|---|---|---|---|---|---|---|
| A | bc1011+bc1031 | bc1011+bc1032 | bc1011+bc1033 | bc1011+bc1034 | bc1011+bc1035 | bc1011+bc1036 | bc1011+bc1037 | bc1011+bc1038 | bc1011+bc1039 | bc1011+bc1040 | bc1019+bc1031 | bc1020+bc1031 |
| B | bc1012+bc1031 | bc1012+bc1032 | bc1012+bc1033 | bc1012+bc1034 | bc1012+bc1035 | bc1012+bc1036 | bc1012+bc1037 | bc1012+bc1038 | bc1012+bc1039 | bc1012+bc1040 | bc1019+bc1032 | bc1020+bc1032 |
| C | bc1013+bc1031 | bc1013+bc1032 | bc1013+bc1033 | bc1013+bc1034 | bc1013+bc1035 | bc1013+bc1036 | bc1013+bc1037 | bc1013+bc1038 | bc1013+bc1039 | bc1013+bc1040 | bc1019+bc1033 | bc1020+bc1033 |
| D | bc1014+bc1031 | bc1014+bc1032 | bc1014+bc1033 | bc1014+bc1034 | bc1014+bc1035 | bc1014+bc1036 | bc1014+bc1037 | bc1014+bc1038 | bc1014+bc1039 | bc1014+bc1040 | bc1019+bc1034 | bc1020+bc1034 |
| E | bc1015+bc1031 | bc1015+bc1032 | bc1015+bc1033 | bc1015+bc1034 | bc1015+bc1035 | bc1015+bc1036 | bc1015+bc1037 | bc1015+bc1038 | bc1015+bc1039 | bc1015+bc1040 | bc1019+bc1035 | bc1020+bc1035 |
| F | bc1016+bc1031 | bc1016+bc1032 | bc1016+bc1033 | bc1016+bc1034 | bc1016+bc1035 | bc1016+bc1036 | bc1016+bc1037 | bc1016+bc1038 | bc1016+bc1039 | bc1016+bc1040 | bc1019+bc1036 | bc1020+bc1036 |
| G | bc1017+bc1031 | bc1017+bc1032 | bc1017+bc1033 | bc1017+bc1034 | bc1017+bc1035 | bc1017+bc1036 | bc1017+bc1037 | bc1017+bc1038 | bc1017+bc1039 | bc1017+bc1040 | bc1019+bc1037 | bc1020+bc1037 |
| H | bc1018+bc1031 | bc1018+bc1032 | bc1018+bc1033 | bc1018+bc1034 | bc1018+bc1035 | bc1018+bc1036 | bc1018+bc1037 | bc1018+bc1038 | bc1018+bc1039 | bc1018+bc1040 | bc1019+bc1038 | bc1020+bc1038 |

**Plate 3**

| | 1 | 2 | 3 | 4 | 5 | 6 | 7 | 8 | 9 | 10 | 11 | 12 |
|---|---|---|---|---|---|---|---|---|---|---|---|---|
| A | bc1001+bc1041 | bc1001+bc1042 | bc1001+bc1043 | bc1001+bc1044 | bc1001+bc1045 | bc1001+bc1046 | bc1001+bc1047 | bc1001+bc1048 | bc1001+bc1049 | bc1001+bc1050 | bc1009+bc1041 | bc1010+bc1041 |
| B | bc1002+bc1041 | bc1002+bc1042 | bc1002+bc1043 | bc1002+bc1044 | bc1002+bc1045 | bc1002+bc1046 | bc1002+bc1047 | bc1002+bc1048 | bc1002+bc1049 | bc1002+bc1050 | bc1009+bc1042 | bc1010+bc1042 |
| C | bc1003+bc1041 | bc1003+bc1042 | bc1003+bc1043 | bc1003+bc1044 | bc1003+bc1045 | bc1003+bc1046 | bc1003+bc1047 | bc1003+bc1048 | bc1003+bc1049 | bc1003+bc1050 | bc1009+bc1043 | bc1010+bc1043 |
| D | bc1004+bc1041 | bc1004+bc1042 | bc1004+bc1043 | bc1004+bc1044 | bc1004+bc1045 | bc1004+bc1046 | bc1004+bc1047 | bc1004+bc1048 | bc1004+bc1049 | bc1004+bc1050 | bc1009+bc1044 | bc1010+bc1044 |
| E | bc1005+bc1041 | bc1005+bc1042 | bc1005+bc1043 | bc1005+bc1044 | bc1005+bc1045 | bc1005+bc1046 | bc1005+bc1047 | bc1005+bc1048 | bc1005+bc1049 | bc1005+bc1050 | bc1009+bc1045 | bc1010+bc1045 |
| F | bc1006+bc1041 | bc1006+bc1042 | bc1006+bc1043 | bc1006+bc1044 | bc1006+bc1045 | bc1006+bc1046 | bc1006+bc1047 | bc1006+bc1048 | bc1006+bc1049 | bc1006+bc1050 | bc1009+bc1046 | bc1010+bc1046 |
| G | bc1007+bc1041 | bc1007+bc1042 | bc1007+bc1043 | bc1007+bc1044 | bc1007+bc1045 | bc1007+bc1046 | bc1007+bc1047 | bc1007+bc1048 | bc1007+bc1049 | bc1007+bc1050 | bc1009+bc1047 | bc1010+bc1047 |
| H | bc1008+bc1041 | bc1008+bc1042 | bc1008+bc1043 | bc1008+bc1044 | bc1008+bc1045 | bc1008+bc1046 | bc1008+bc1047 | bc1008+bc1048 | bc1008+bc1049 | bc1008+bc1050 | bc1009+bc1048 | bc1010+bc1048 |

**Plate 4**

| | 1 | 2 | 3 | 4 | 5 | 6 | 7 | 8 | 9 | 10 | 11 | 12 |
|---|---|---|---|---|---|---|---|---|---|---|---|---|
| A | bc1011+bc1041 | bc1011+bc1042 | bc1011+bc1043 | bc1011+bc1044 | bc1011+bc1045 | bc1011+bc1046 | bc1011+bc1047 | bc1011+bc1048 | bc1011+bc1049 | bc1011+bc1050 | bc1019+bc1041 | bc1020+bc1041 |
| B | bc1012+bc1041 | bc1012+bc1042 | bc1012+bc1043 | bc1012+bc1044 | bc1012+bc1045 | bc1012+bc1046 | bc1012+bc1047 | bc1012+bc1048 | bc1012+bc1049 | bc1012+bc1050 | bc1019+bc1042 | bc1020+bc1042 |
| C | bc1013+bc1041 | bc1013+bc1042 | bc1013+bc1043 | bc1013+bc1044 | bc1013+bc1045 | bc1013+bc1046 | bc1013+bc1047 | bc1013+bc1048 | bc1013+bc1049 | bc1013+bc1050 | bc1019+bc1043 | bc1020+bc1043 |
| D | bc1014+bc1041 | bc1014+bc1042 | bc1014+bc1043 | bc1014+bc1044 | bc1014+bc1045 | bc1014+bc1046 | bc1014+bc1047 | bc1014+bc1048 | bc1014+bc1049 | bc1014+bc1050 | bc1019+bc1044 | bc1020+bc1044 |
| E | bc1015+bc1041 | bc1015+bc1042 | bc1015+bc1043 | bc1015+bc1044 | bc1015+bc1045 | bc1015+bc1046 | bc1015+bc1047 | bc1015+bc1048 | bc1015+bc1049 | bc1015+bc1050 | bc1019+bc1045 | bc1020+bc1045 |
| F | bc1016+bc1041 | bc1016+bc1042 | bc1016+bc1043 | bc1016+bc1044 | bc1016+bc1045 | bc1016+bc1046 | bc1016+bc1047 | bc1016+bc1048 | bc1016+bc1049 | bc1016+bc1050 | bc1019+bc1046 | bc1020+bc1046 |
| G | bc1017+bc1041 | bc1017+bc1042 | bc1017+bc1043 | bc1017+bc1044 | bc1017+bc1045 | bc1017+bc1046 | bc1017+bc1047 | bc1017+bc1048 | bc1017+bc1049 | bc1017+bc1050 | bc1019+bc1047 | bc1020+bc1047 |
| H | bc1018+bc1041 | bc1018+bc1042 | bc1018+bc1043 | bc1018+bc1044 | bc1018+bc1045 | bc1018+bc1046 | bc1018+bc1047 | bc1018+bc1048 | bc1018+bc1049 | bc1018+bc1050 | bc1019+bc1048 | bc1020+bc1048 |

**Supplementary Fig. 9.** Scheme of asymmetrical barcoding for a 384-plex genotyping configuration

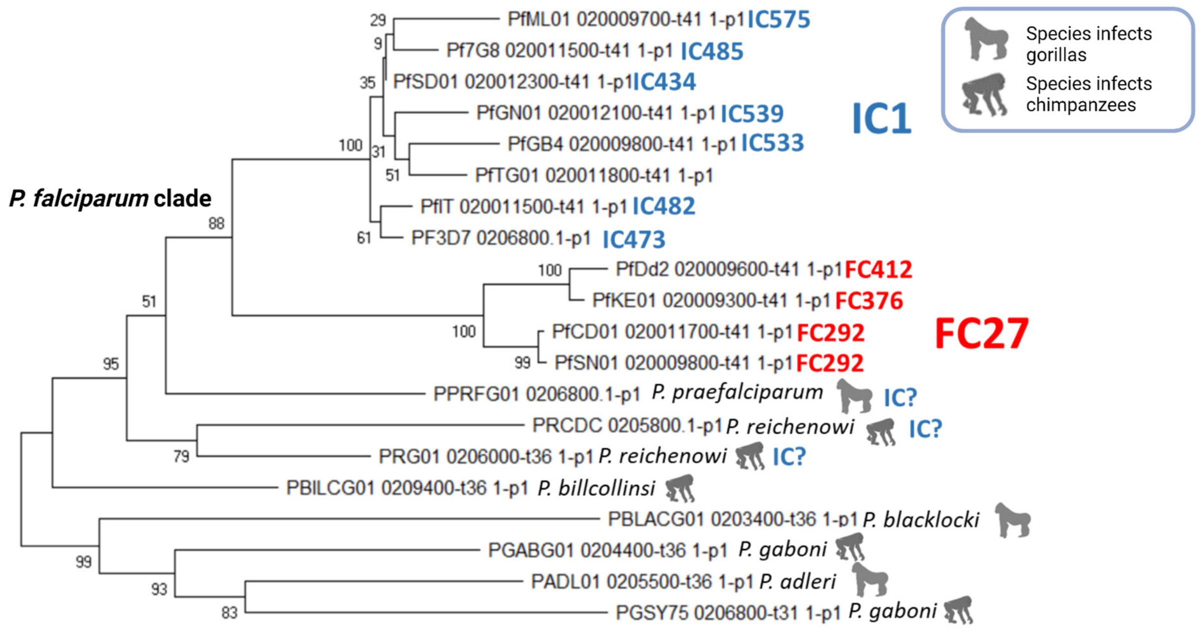


**Supplementary Fig. 10. Phylogeny for reference *msp2* orthologs in the subgenus *Laverania*.** Neighbor-joining method was used for the construction of the tree. 1000 bootstrapping iterations were run; numbers indicate the percentage of trees supporting each branch. The ortholog of *msp2* in *P. billcollinsi* is the only one from the subgenus *Laverania* that is closely related to the *P. falciparum* clade and where no subfamily (IC1 or FC27) can be identified. Sequence IDs are shown at the branch tips.
